# HLJ1 amplifies endotoxin–induced sepsis severity by promoting IL-12 heterodimerization in macrophages

**DOI:** 10.1101/2022.01.05.475083

**Authors:** Wei-Jia Luo, Sung-Liang Yu, Chia-Ching Chang, Min-Hui Chien, Keng-Mao Liao, Ya-Ling Chang, Ya-Hui Chuang, Jeremy J.W. Chen, Pan-Chyr Yang, Kang-Yi Su

## Abstract

Heat shock protein (HSP) 40 has emerged as a key actor in both innate and adaptive immunity, whereas the role of HLJ1, a molecular chaperone in HSP40 family, in modulating endotoxin–induced sepsis severity is still unclear. Here, we use single-cell RNA sequencing to characterize mouse liver nonparenchymal cell populations under LPS (lipopolysaccharide) stimulation, and show that HLJ1 deletion affected IFN-γ-related gene signatures in distinct immune cell clusters. HLJ1 deficiency also leads to reduced serum levels of IL-12 in LPS-treated mice, contributing to dampened production of IFN-γ in natural killer cells but not CD4^+^ or CD8^+^ T cells, and subsequently to improved survival rate. Adoptive transfer of HLJ1-deleted macrophages into LPS-treated mice results in reduced IL-12 and IFN-γ levels and protects the mice from IFN-γ–dependent mortality. In the context of molecular mechanisms, HLJ1 is an LPS-inducible protein in macrophages and converts misfolded IL-12p35 homodimers to monomers, which maintains bioactive IL-12p70 heterodimerization and secretion. This study suggests HLJ1 causes IFN-γ–dependent septic lethality by promoting IL-12 heterodimerization, and targeting HLJ1 has therapeutic potential in inflammatory diseases involving activating IL-12/IFN-γ axis.

## Introduction

Sepsis, causes systemic inflammatory response syndrome, is a complex disorder defined as life-threatening organ dysfunction that occurs as a result of a dysregulated host response to pathogen infection (Singer et al., 2016). In high-income countries, sepsis remains a major health problem, since it has an in-hospital mortality rate of 21.2%, and approximately 2.8 million deaths per year occur as a result of multi-organ failure related to sepsis (Adhikari et al., 2010; Vincent et al., 2014). Sepsis is a biphasic disorder characterized by an initial inflammatory phase followed by a prolonged immunosuppressive phase (Hotchkiss et al., 2013). In fact, in COVID-19 patients, sepsis comprises up to 60% of complications (Zhou et al., 2020). The main treatments for sepsis are antibiotic administration and supportive therapies, although precision medicine-based strategies and markers for patient selection are still under investigation (Angus & van der Poll, 2013; Pierrakos et al., 2020). Cytokine storm, which can be triggered by pathogen infection, is a life-threatening systemic inflammatory response featuring elevated levels of circulating cytokines and immune cell hyperactivation, which can cause severe sepsis and can dramatically increase mortality (Chousterman et al., 2017; Fajgenbaum & June, 2020). When the host is exposed to bacterial endotoxins, macrophages initiate an inflammatory response by sensing microbial products via the TLRs expressed in their cell membranes or within their endosomes (Heymann & Tacke, 2016; Mencin et al., 2009). TLR signaling plays a critical role in generating sepsis-associated cytokine storm, which can cause severe systemic inflammatory conditions such as multiple organ dysfunction syndrome and even sepsis death (Kumar, 2020). For example, LPS-mediated TLR4 signaling has been shown to trigger inflammatory responses that contribute to various liver diseases in both human and mouse models (Jirillo et al., 2002; Mencin et al., 2009; Rivera et al., 2007). Over the past decade, understanding of the molecular mechanisms responsible for the initial recognition of LPS and the ultimate generation of cytokine storm has increased dramatically. LPS binds to an LPS-binding protein and then forms a complex with myeloid differentiation 2 and cluster of differentiation 14, which is recognized by TLR4 (Yang et al., 2000). The engagement of TLR4 transduces signals to intracellular proteins, resulting in a cascade of molecular responses and the final production of various inflammatory cytokines (Beutler & Rietschel, 2003). Despite abundant studies, little is known about the precise pathogenesis and immune modulation of sepsis (Angus & van der Poll, 2013).

Heat shock proteins (HSPs) can be upregulated under certain cellular-stress conditions, such as oxidative stress, hypoxia, fever, and inflammation, and they act as chaperones to maintain the functions of cytosolic proteins (Georgopoulos & Welch, 1993; Rosenzweig et al., 2019). Studies have implicated serum HSP70 as a biomarker for inflammatory processes in multiple sclerosis and shown that, in sepsis, serum concentrations of HSP70 are elevated, which is associated with increased mortality (Gelain et al., 2011; Lechner et al., 2018). HSP40 is known to induce proinflammatory cytokine production in macrophages and can activate bone marrow-derived dendritic cells through recognition of TLR4, leading to subsequent activation of MAPK, NF-κB, and PI3K–Akt pathways (Cui et al., 2017; Wu et al., 2017). In the context of rheumatoid arthritis, a study has demonstrated that HSP40 inhibits T cell division and stimulates anti-inflammatory cytokine production mediated by peripheral blood mononuclear cells (PBMCs) (Tukaj et al., 2010). Even though knowledge about the immunomodulatory role of HSPs is growing, how they mediate endotoxin-induced sepsis and the underlying mechanisms remain unclear.

Human liver DnaJ-like protein (HLJ1) is a member of heat shock protein 40 family (HSP40) and is also known as DNAJB4. In humans, HLJ1 is considered to be a tumor suppressor in non-small-cell lung cancer and colorectal cancer, since its upregulation suppresses tumor invasion and high expression of HLJ1 is associated with prolonged survival of patients (Liu et al., 2014; Tsai et al., 2006; Wang et al., 2005). However, the actual immunomodulatory role of HLJ1 in sepsis remains unclear. Recently, chaperones have emerged as mediators of IL-12 family protein folding, assembly, and degradation (Meier et al., 2019; Reitberger et al., 2017). DnaJ HSP40 member C10 (DNAJC10), also known as ERdj5, mediates endoplasmic reticulum-associated degradation (ERAD) by reducing incorrect disulfide bonding in misfolded glycoproteins recognized by EDEM1 (Ushioda et al., 2008). Bioactive IL-12 is a disulfide-bridged heterodimeric glycoprotein that consists mainly of an α subunit (IL-12p35) and a β subunit (IL-12p40) (Yoon et al., 2000). ERdj5 has been shown to reduce disulfide bonding in non-native homodimer IL-12p35 to low-molecular-weight (LMW) IL-12p35 monomers (Reitberger et al., 2017). Despite a growing understanding of chaperone-mediated IL-12 family protein folding and assembly, the precise mechanisms by which HLJ1 regulates IL-12 biosynthesis and the subsequent immune response have not yet been elucidated.

To further elucidate the mechanisms by which HLJ1 governs immune responses during sepsis, we determined the immune profile that is affected by HLJ1 under LPS stress. With single-cell RNA sequencing analysis, we found an impaired IFN-γ-related gene signature in murine nonparenchymal liver cells in which HLJ1 had been deleted. LPS-inducible HLJ1 reduced the accumulation of misfolded IL-12p35 homodimer and enhanced IL-12p70 dimerization in macrophages, which led to augmented IFN-γ production, mainly by hepatic and splenic natural killer (NK) cells, contributing to sepsis-related mortality. The current study reveals the previously unknown role of HLJ1 in promoting IFN-γ–dependent endotoxic death via the regulation of IL-12 folding and biosynthesis in macrophages. Therefore, targeting HLJ1 provides a strategy for developing therapeutic approaches to inflammatory diseases involving the activation of the IL-12/IFN-γ axis.

## Results

### HLJ1 deficiency protected mice against lethal septic shock

To address the question of whether HLJ1 participates in regulating LPS-induced systemic inflammatory responses, HLJ1-deficient (*Hlj1^−/−^*) mice and wild-type littermates (*Hlj1^+/+^*) were intraperitoneally injected with LPS derived from Gram-negative bacteria. *Hlj1^+/+^* and *Hlj1^−/−^* mice showed similar survival rates when low-dose LPS (10 mg/kg) was administered (Figure 1A, right panel), but *Hlj1^−/−^* mice were significantly more resistant to LPS-induced sepsis and exhibited longer survival than *Hlj1^+/+^* mice when subjected to a higher lethal dose of LPS (20 mg/kg) (Figure 1A, left panel). Since LPS is known to induce systemic immune responses, we analyzed complete blood counts (CBCs) of peripheral blood and pathological changes in the mice injected with LPS. The two genotypes had similar counts and percentages of white blood cells, neutrophils, lymphocytes, monocytes, and eosinophils at 4 h and 8 h post-LPS injection (Figure 1 – figure supplement 1A). Although LPS is known to cause liver dysfunction and damage, there was no difference in serum aspartate transaminase (ALT) levels, an indicator of liver damage, between the two genotypes (Figure 1 – figure supplement 1B). Since LPS possesses a high affinity for high-density lipoprotein (HDL) and low-density lipoprotein (LDL), causing it to be carried in the circulation and transported to the liver, we also examined serum levels of HDL and LDL. The quantities of both lipoproteins were slightly reduced 4 h post-injection, but there was no difference between the two genotypes (Figure 1 – figure supplement 1C).

**Figure 1.**
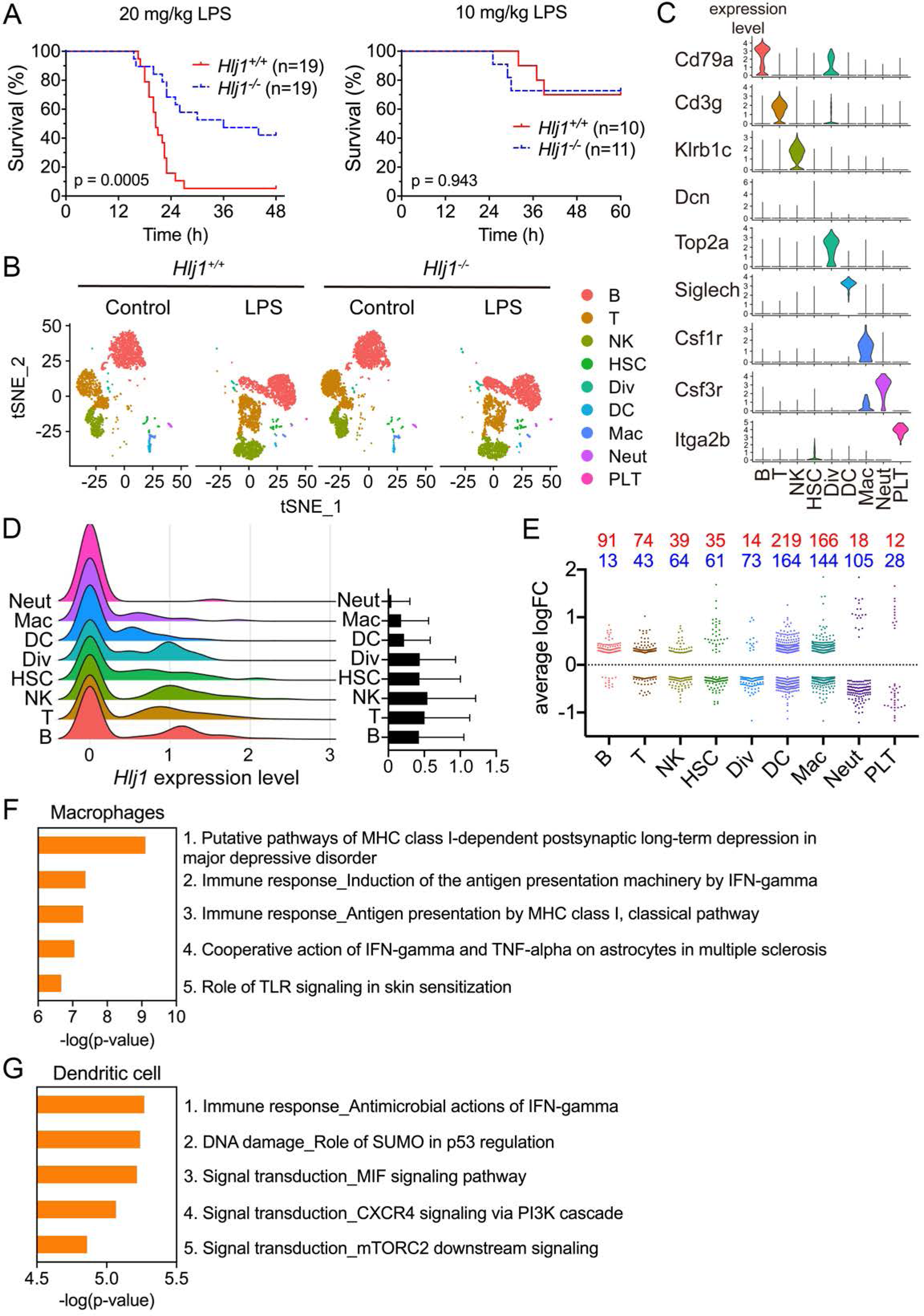
*Hlj1^−/−^* mice survive better than *Hlj1^+/+^* mice after high-dose LPS injection. (**A**) Kaplan–Meier analysis of the overall survival of 6–8-week-old *Hlj1^+/+^* mice and *Hlj1^−/−^* mice injected with low-dose (10 mg/kg, n = 10–11) or high-dose (20 mg/kg, n = 19) LPS. (**B**) Mice were injected with 20 mg/kg LPS, or PBS as a control, and after 8 h hepatic nonparenchymal cells were isolated for scRNA-seq analysis. The plot shows the t-SNE visualization of liver nonparenchymal cell clusters based on 11,651 single-cell transcriptomes. B, B cells; T, T cells; NK, NK cells; HSC, hepatic stellate cells; Div, dividing cells; DC, dendritic cells; Mac, macrophages; Neut, neutrophils; PLT, platelets. (**C**) Expression levels of representative known marker genes for each cluster. (**D**) Visualization of expression distribution of the *Hlj1* gene in each cluster of cells in PBS-treated *Hlj1^+/+^* mice. Data presented are means ± SD. (**E**) Cell-type distribution and log-transformed expression fold change (logFC) for upregulated (red) and downregulated (blue) genes from a comparison of LPS-treated *Hlj1^+/+^* mice with LPS-treated *Hlj1^−/−^* mice. The Wilcoxon rank–sum test was used to identify differentially expressed genes (*P* < 0.05, |logFC| > 0.25). (**F** and **G**) Enrichment analysis showing ranked pathway signatures associated with up- and downregulated genes (P < 0.05, |logFC| > 0.25) from a comparison of macrophages (F) and dendritic cells (G) from LPS-injected *Hlj1^+/+^* mice with those from *Hlj1^−/−^* mice. **Source data 1.** Data for graphs depicted in Figure 1A, E-G.

### HLJ1 deficiency resulted in reduced IFN-γ-related signatures in single-cell RNA sequencing analysis

To comprehensively identify HLJ1 as a potential immune modulator, as well as to understand the underlying mechanism by which HLJ1 regulates LPS-induced immune responses in the liver, we performed a single-cell RNA sequencing (scRNA-seq) analysis of hepatic nonparenchymal cells. We acquired a T-distributed stochastic neighbor embedding (t-SNE) map of 11,651 single-cell transcriptomes from the livers of *Hlj1^+/+^* and *Hlj1^−/−^* mice injected with either LPS or saline (Figure 1B). Since apoptotic mammalian cells express mitochondrial genes and export the gene transcripts to the cytoplasm, we performed quality control to exclude cells with high levels of mtDNA expression and retain only high-quality cells (Figure 1 – figure supplement 2A). We also excluded cells with excessive unique molecular identifier (UMI) counts and genes (Figure 1 – figure supplement 2A). Distinct clusters on the t-SNE visualization revealed nine cell types that were identified based on well-known marker genes published in previous studies (Xiong et al., 2019; Zhao et al., 2019) (Figure 1C, Figure 1 – figure supplement 2B and C). LPS stimulation significantly altered gene-expression and cell clustering patterns in both genotypes (Figure 1B). HLJ1 was abundant in all cell types, without significant differences among clusters (Figure 1D). We analyzed differentially expressed genes with absolute log-fold changes greater than 0.25 and P-values less than 0.05 after LPS induction based on their cell type. In LPS-treated mice, HLJ1 deletion led to most genes being significantly upregulated in dendritic cells, followed by macrophages, and then B cells; on the other hand, downregulated genes were mainly found in macrophages, dendritic cells, and neutrophils (Figure 1E). We therefore selected genes with significant changes in macrophages and dendritic cells for pathway analysis.

Enrichment analysis of these significant genes from LPS-injected *Hlj1^+/+^* and *Hlj1^−/−^* mice revealed differential expression not only in IFN-γ-activating pathways, but also in MHC class-I–related signals in macrophages (Figure 1F). In dendritic cells, the differentially expressed genes were mainly enriched in IFN-γ-stimulating immune-response pathways and macrophage migration inhibitory factor (MIF) signaling pathways (Figure 1G). Hence, we focused on IFN-γ expression levels in individual cells and found that there were indeed significantly fewer IFN-γ–positive cells in LPS-injected *Hlj1^−/−^* mice than *Hlj1^+/+^* mice (Figure 2A). The extent of IFN-γ induction depended on cell type: LPS treatment led to significantly elevated IFN-γ transcription in NK, T, and B cells, but not in other cell types (Figure 2B). IFN-γ expression patterns at the single-cell level among NK, T, and B cells indicated specific distinct clusters of IFN-γ–positive cells in the livers of LPS-injected mice compared to those of control mice (Figure 2C). T and B cells in LPS-treated *Hlj1^+/+^* and *Hlj1^−/−^* mice exhibited comparable levels of IFN-γ, but the number of IFN-γ–positive NK cells was lower in *Hlj1^−/−^* mice (Figure 2C). Indeed, process network analysis showed that significantly differentially expressed genes from the NK cells of LPS-treated *Hlj1^+/+^* and *Hlj1^−/−^* mice were mainly enriched in inflammatory pathways involving IFN-γ signaling and IFN-γ–related NK cell cytotoxicity (Figure 2D). To further validate the results from the scRNA-seq analysis, we analyzed hepatic IFN-γ expression levels via quantitative real-time PCR (qRT-PCR). We found that there were lower levels of IFN-γ transcripts in *Hlj1^−/−^* mice than in *Hlj1^+/+^* mice 4 h and 8 h after LPS injection (Figure 2E). These results suggested that HLJ1 plays an important role in promoting severe systemic immune responses via the enhancement of IFN-γ production mediated by NK cells and the alteration of the IFN-γ–related gene signature in endotoxin-induced sepsis.

**Figure 2.**
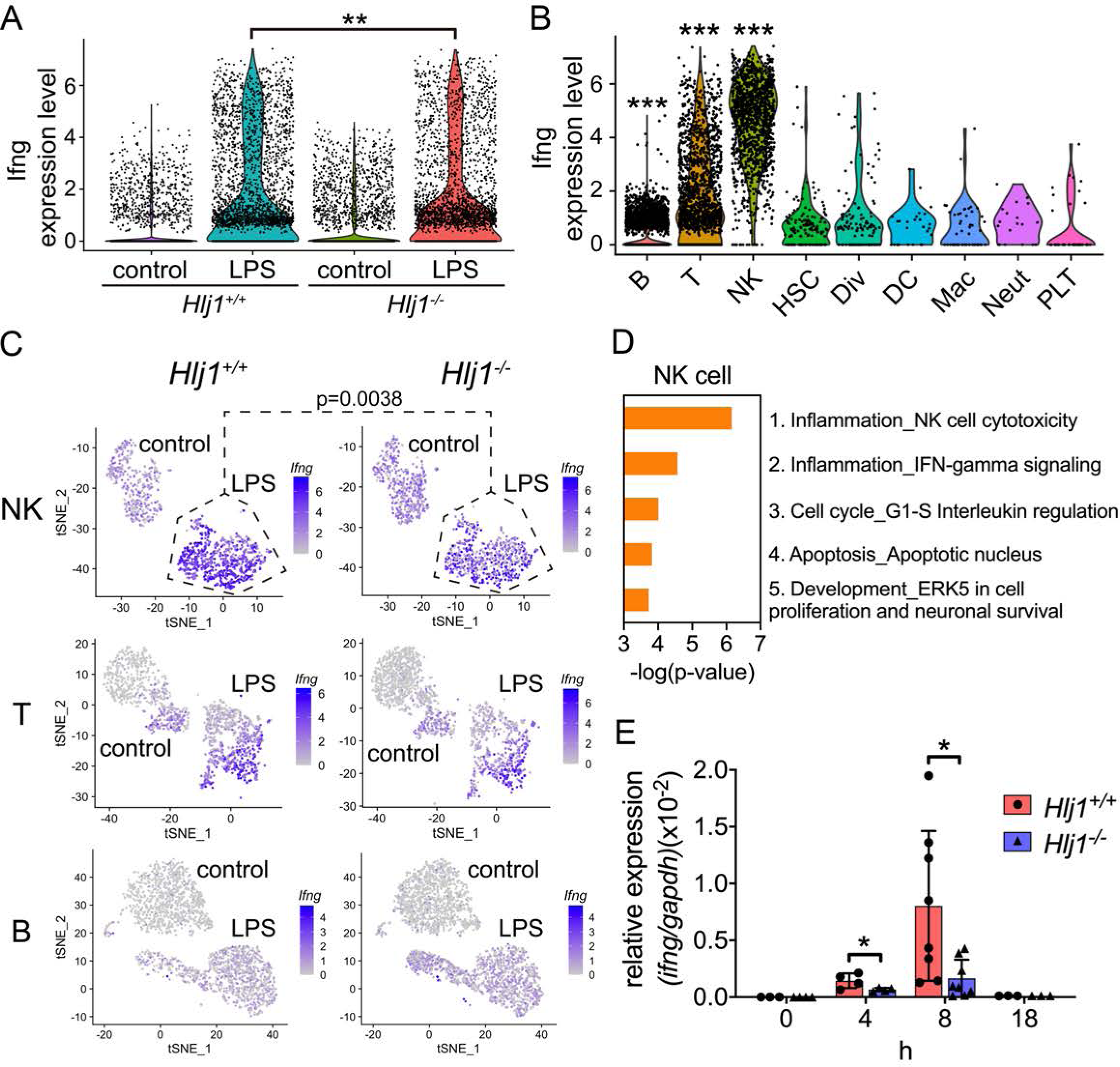
HLJ1 deficiency leads to altered IFN-γ-related signatures in NK cells under LPS stress. (**A**) IFN-γ gene expression levels in each cell. Significance was calculated using the Wilcoxon rank–sum test; *P* = 0.009. (**B**) IFN-γ expression patterns in B (*P* < 0.001), T (*P* < 0.001), and NK (*P* < 0.001) cells and other clusters in LPS-treated mice. Significance was calculated using the Wilcoxon rank–sum test. (**C**) t-SNE visualization of IFN-γ expression profiles in NK, T, and B cells isolated from *Hlj1^+/+^* and *Hlj1^−/−^* mice injected with LPS. (**D**) Enrichment analysis showing ranked network signatures associated with up- and downregulated genes (*P* < 0.05, |logFC| > 0.25) from a comparison of NK cells from LPS-injected *Hlj1^+/+^* mice with NK cells from *Hlj1^−/−^* mice. (**E**) *Hlj1^+/+^* and *Hlj1^−/−^* mice were injected with LPS and, at the indicated time points, whole liver mRNA was extracted for the measurement of hepatic IFN-γ expression levels via qRT-PCR. *P* = 0.026 and *P* = 0.014 for the 4 h and 8 h groups, respectively. Data presented are means ± SD. * *P* < 0.05, ** *P* < 0.01, *** *P* < 0.001. **Source data 1.** Data for graphs depicted in Figure 2D and E.

### HLJ1 deletion alleviates IFN-γ–dependent septic death

Cytokine storm caused by a dysregulated immune response to infection is the major cause of septic shock and multiple organ failure, and can contribute to sepsis-associated death. It is thus important to quantify cytokine levels after LPS treatment. We used a bead-based immunoassay to determine serum levels of multiple LPS-induced cytokines and chemokines. Among the assayed cytokines, IL-1α and IL-12p70 levels were significantly and slightly lower, respectively, in *Hlj1^−/−^* mice (Figure 3A). Notably, serum IFN-γ levels were also significantly lower, which was consistent with the results obtained in the scRNA-seq analysis (Figure 3A). We further confirmed these results via ELISA and found that, indeed, serum IFN-γ levels were significantly lower in *Hlj1^−/−^* mice than in *Hlj1^+/+^* mice 8 h and 18 h after LPS treatment (Figure 3B), whereas there was no difference in IL-1α between the two genotypes 8 h after LPS induction (Figure 3C). The effect of HLJ1 deletion on IFN-γ production was also demonstrated by using a lower dose of LPS (4 mg/kg) which was able to cause moderate endotoxemia. In line with the effects found during severe endotoxemia, HLJ1 deletion led to reduced serum levels of IFN-γ when mice were challenged with a non-lethal dose of LPS (Figure 3D). To confirm that the mitigation of LPS-induced septic death in *Hlj1^−/−^* mice was not due to a change in their susceptibility to cytokine storm, we analyzed the correlation between serum levels of IFN-γ and survival status (Figure 3E). As it turned out, mice bearing high serum levels of IFN-γ died, while those with low levels survived, regardless of genotype, suggesting that HLJ1 deletion does not confer increased susceptibility to IFN-γ. When *Hlj1^+/+^* mice were injected with anti-IFN-γ neutralizing antibodies prior to LPS treatment, they exhibited significantly improved survival (Figure 3F). However, the survival rate of *Hlj1^−/−^* mice was only slightly improved after IFN-γ neutralization. These results indicated that HLJ1 enhances septic death by augmenting IFN-γ signaling.

**Figure 3.**
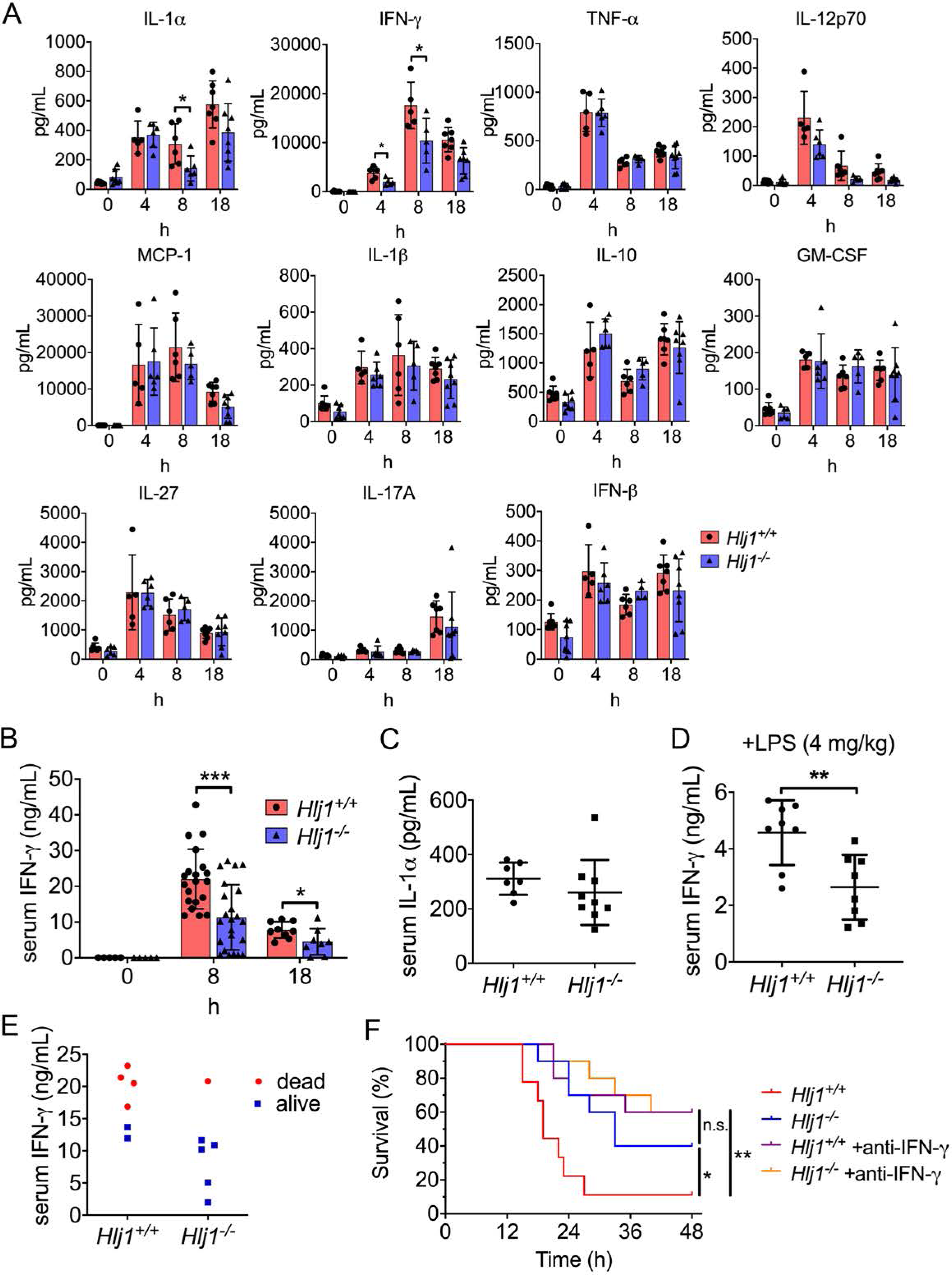
HLJ1 deletion alleviates IFN-γ–dependent cytokine storm and death. **(A)** Serum from *Hlj1^+/+^* and *Hlj1^−/−^* mice administered with LPS was analyzed at the indicated time points to quantify 11 cytokines via a cytokine bead array. IL-1α, *P* = 0.03; 4 h IFN-γ, *P* = 0.027; 8 h IFN-γ, *P* = 0.04; n = 5–8 per group. **(B)** Serum IFN-γ levels were quantified using ELISA 8 h (n = 20–22) and 18 h (n = 8–9) after LPS injection. 8 h IFN-γ, *P* < 0.001; 18 h IFN-γ, *P* = 0.039. **(C)** Serum IL-1α levels from n= 8–9 mice were quantified 8 h after LPS injection. **(D)** Mice (n = 8 biological replicates) were injected with lower dose 4 mg/kg LPS and after 8 hours serum were collected for quantification of IFN-γ levels. *P* = 0.005. **(E)** Correlation between survival status and serum IFN-γ levels in *Hlj1^+/+^* and *Hlj1^−/−^* mice injected with LPS (n = 6 per group). (**F**) Kaplan–Meier analysis of overall survival of *Hlj1^+/+^* and *Hlj1^−/−^* mice (n = 9–10) injected with 100 μg anti-IFN-γ neutralizing antibodies 1 h before LPS (20 mg/kg) challenge. *Hlj1^+/+^* vs *Hlj1^−/−^* mice, *P* = 0.015; *Hlj1^+/+^* vs *Hlj1^+/+^*+anti-IFN-γ, *P* = 0.007. Data presented are means ± SD. * *P* < 0.05, ** *P* < 0.01, *** *P* < 0.001. **Source data 1.** Data for graphs depicted in Figure 3A-F.

### Intracellular IFN-γ decreased in splenic NK cells after HLJ1 deletion

Since LPS has been reported to induce systemic inflammatory responses, we analyzed splenic T, NK, and B cell populations in *Hlj1^+/+^* and *Hlj1^−/−^* mice. The percentages (Figure 4 – figure supplement 1A and B) and sizes (Figure 4 – figure supplement 1C) of splenic CD4^+^, CD8^+^, NK, and B cell populations in LPS-treated *Hlj1^−/−^* mice were similar to those in *Hlj1^+/+^* mice. Our scRNA-seq data showed that IFN-γ was mainly secreted by NK cells in LPS-treated mice (Figure 2B and C), and we accordingly performed intracellular IFN-γ staining with flow cytometry analysis of splenic immune cells. In LPS-treated mice, the percentage of IFN-γ^+^ CD4^+^ and IFN-γ^+^ CD8^+^ T cells was slightly lower in LPS-injected *Hlj1^−/−^* mice than in *Hlj1^+/+^* mice, while that of IFN-γ^+^ NK cells was significantly lower (Figure 4A), which is in accordance with our findings from the scRNA-seq analysis of mouse hepatic cells. Transcriptional levels of IFN-γ were also lower in the spleens of LPS-treated *Hlj1^−/−^* mice than in *Hlj1^+/+^* mice (Figure 4B), which implied weakened upstream signaling leading to reduced IFN-γ levels. We therefore tested IFN-γ expression in NK cells from *Hlj1^−/−^* and *Hlj1^+/+^* mice under IL-12 stimulation. We isolated primary NK cells from spleens from both genotypes with ∼90% purity (Figure 4C), then stimulated the cells with recombinant mouse IL-12p70 for 24 h, and finally quantified the supernatant IFN-γ via ELISA (Figure 4D). This indicated that IFN-γ expression was induced in an IL-12p70-dependent manner, and the amount of supernatant IFN-γ produced by HLJ1-deleted NK cells was comparable to that produced by wild-type NK cells in response to IL-12p70 stimulation (Figure 4D). Combined, these results suggested that *Hlj1* deletion leads to reduced IFN-γ production in not only hepatic but also splenic NK cells, whereas HLJ1 deficiency alters neither the sensitivity to IL-12p70 nor the ability to secrete IFN-γ of NK cells.

**Figure 4.**
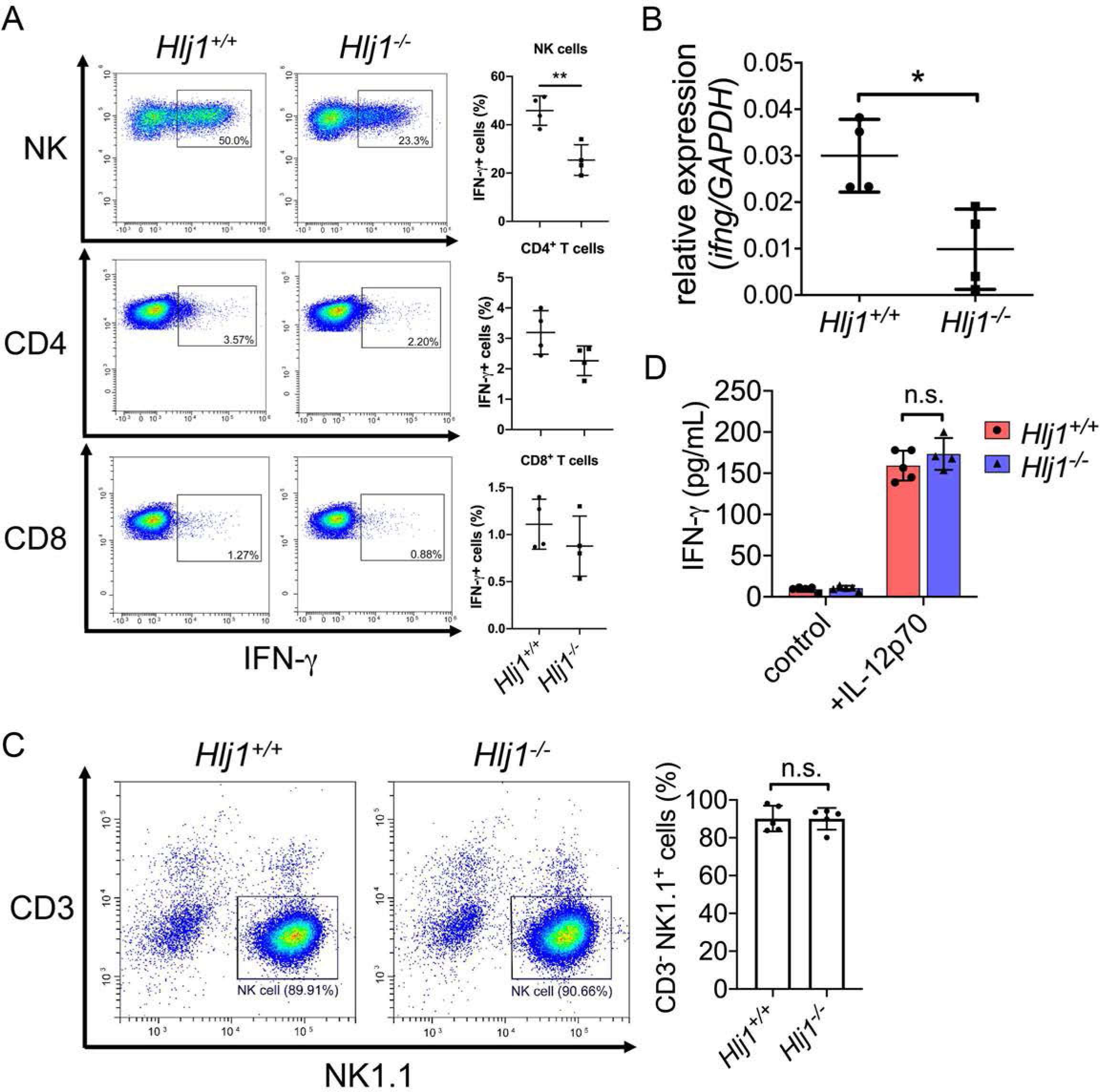
Intracellular IFN-γ levels decreased in splenic NK cells after HLJ1 deletion. (**A)** *Hlj1^+/+^* and *Hlj1^−/−^* mice (n = 4 per group) were injected intraperitoneally with 20 mg/kg LPS, and splenocytes were isolated after 2.5 h. Expression of intracellular IFN-γ levels in *Hlj1^+/+^* and *Hlj1^−/−^* NK, CD4^+^ T, and CD8^+^ T cells were detected via flow cytometry analysis. NK cells, *P* = 0.004. Representative samples are shown. (**B**) RNA from n = 4 mice spleens were isolated 4 h after LPS administration, and transcriptional levels of IFN-γ were quantified via qRT-PCR; *P* = 0.014. (**C**) Expression of NK1.1 in primary NK cells isolated from *Hlj1^+/+^* and *Hlj1^−/−^* mice (n = 5 per group) was detected via flow cytometry. Representative samples are shown. (**D**) Primary NK cells purified from *Hlj1^+/+^* and *Hlj1^−/−^* mice spleens were treated with 10 ng/mL IL-12p70 for 24 h and supernatant IFN-γ was quantified using ELISA (n = 4–5 biological replicates). Data presented are means ± SD. * *P* < 0.05, ** *P* < 0.01, n.s., not significant. **Source data 1.** Data for graphs depicted in Figure 4A-D.

### HLJ1 contributes to IL-12–dependent IFN-γ cytokine storm and lethality

Kupffer cells are the macrophages of the liver, and they are responsible for clearing bacteria and endotoxins from the blood stream. In response to LPS stimulation, liver-resident Kupffer cells release proinflammatory cytokines and nitric oxide to initiate the inflammatory cascade. We therefore examined the number of Kupffer cells and liver mononuclear cells via F4/80 fluorescent staining, which showed no significant difference between *Hlj1^−/−^* and *Hlj1^+/+^* mice (Figure 5 – figure supplement 1A and B). This suggested that HLJ1 deficiency has little impact on macrophage infiltration before and after LPS administration. Since LPS-induced inflammatory liver injury features activation of the NF-κB pathway and the production of multiple inflammatory cytokines for subsequent IFN-γ activation, we next focused on the cytokines and chemokines generated by hepatic macrophages after LPS stimulation. Transcription levels of proinflammatory cytokines IL-1β, TNF-α, and monocyte chemoattractant protein-1 (MCP-1) did not differ significantly between genotypes, but IL-6 was downregulated in *Hlj1^−/−^* livers compared to *Hlj1^+/+^* livers (Figure 5A). In addition, transcriptional levels of hepatic IL-12, a cytokine contributing to LPS-induced septic death via cytokine storm mediated by the IL-12/18–IFN-γ axis, were lower in *Hlj1^−/−^* than in *Hlj1^+/+^* mice, although IL-18 expression levels were similar in the two genotypes (Figure 5B). Intriguingly, serum levels of IL-12p70 but not IL-6 were dramatically decreased in HLJ1–deficient mice (Figure 5C). To investigate the impact of IL-12 on IFN-γ production in the context of HLJ1 deficiency, we administered anti-IL-12 neutralizing antibodies intraperitoneally 1 h before the LPS challenge. The serum IL-12 induced by LPS was efficiently neutralized by the antibodies, and serum IFN-γ was also dramatically reduced in both genotypes (Figure 5D). It is noteworthy that the serum levels of IFN-γ in anti–IL-12 antibody–injected *Hlj1^−/−^* mice were comparable to those in *Hlj1^+/+^* mice during sepsis (Figure 5D). More importantly, IL-12 neutralization significantly reduced the mortality of *Hlj1^+/+^* mice, which displayed similar survival to *Hlj1^−/−^* mice when challenged with LPS (Figure 5E). Combined, these results suggested that HLJ1 is required for LPS-induced IL-12 production, subsequent IFN-γ release, and eventual sepsis death.

**Figure 5.**
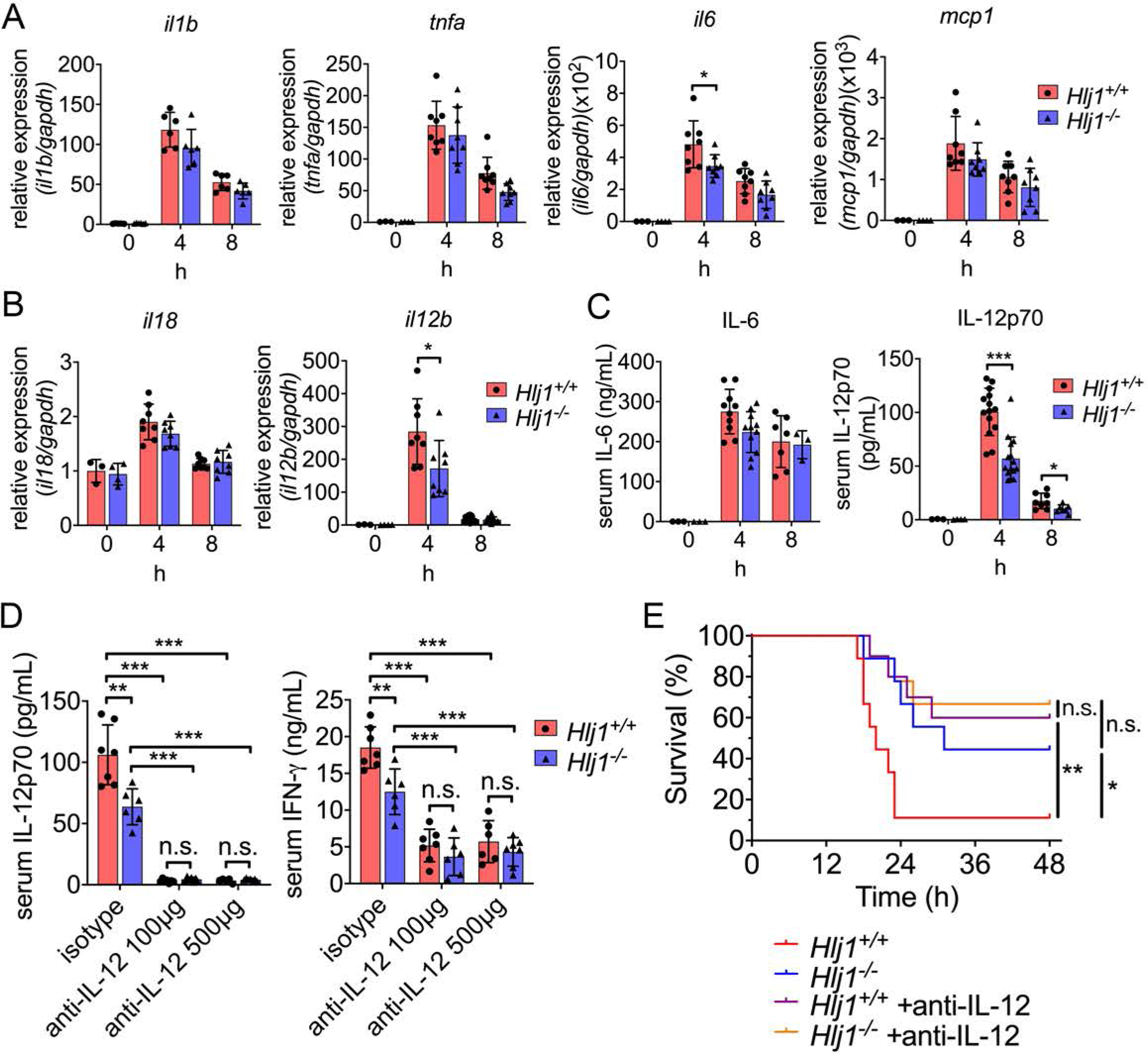
HLJ1 deletion alleviates IL-12–dependent septic death. *Hlj1^+/+^* and *Hlj1^−/−^* mice were intraperitoneally injected with 20 mg/kg LPS. (**A** and **B**) After 4 h or 8 h, the RNA from n = 6–8 total livers was isolated and gene expression levels were quantified via qRT-PCR. IL-6, *P* = 0.033; IL-12b, *P* = 0.029. (**C**) Serum levels of IL-12p70 and IL-6 in LPS-treated *Hlj1^+/+^* and *Hlj1^−/−^* mice were quantified via ELISA 4 h (n = 11–14) and 8 h (n = 4–7) after LPS administration. 4 h IL-12p70, *P* < 0.001; 8 h IL-12p70, *P* = 0.033. (**D**) *Hlj1^+/+^* and *Hlj1^−/−^* mice were intraperitoneally injected with anti-IL-12 neutralizing antibodies 1 h prior to the injection of 20 mg/kg LPS. After the administration of LPS and anti-IL-12 antibodies, the serum was collected at the indicated time points and analyzed for IL-12 and IFN-γ levels. (**E**) Kaplan–Meier analysis of overall survival of *Hlj1^+/+^* and *Hlj1^−/−^* mice injected with 100 μg anti-IL-12 neutralizing antibodies 1 h before the 20 mg/kg LPS challenge (n = 9–11 per group). *Hlj1^+/+^* vs *Hlj1^−/−^* mice, *P* = 0.014; *Hlj1^+/+^* vs *Hlj1^+/+^*+anti-IFN-γ, *P* = 0.007. Data presented are means ± SD. * *P* < 0.05, ** *P* < 0.01, *** *P* < 0.001, n.s., not significant. **Source data 1.** Data for graphs depicted in Figure 5A-E.

### HLJ1 functions in macrophages to maintain the IL-12/IFN-γ axis *in vivo* and promote septic death

To confirm that HLJ1 functions exclusively in macrophages, leading to LPS-induced IL-12 secretion, IFN-γ–related cytokine storm, and septic death, we performed macrophage transplantation. This was achieved by depleting macrophages and Kupffer cells with clodronate liposomes, followed by adoptive transfer of macrophages from other mice. We intravenously transplanted bone marrow-derived macrophages (BMDMs) into the mice 48 h after the intravenous injection of clodronate liposomes. This was followed by LPS treatment at 72 h and blood sampling at 76 h and 80 h (Figure 6A). The administration of clodronate liposomes efficiently depleted most of the liver-resident macrophages in both mice genotypes, since we observed few F4/80-positive cells after this process (Figure 6B). Serum levels of IL-12 and IFN-γ in the LPS-treated mice were significantly reduced as a result of this depletion (Figure 6C). After the depletion, the LPS-treated *Hlj1^−/−^* mice into which *Hlj1^+/+^* BMDMs had been transplanted exhibited higher serum levels of IL-12 and IFN-γ than those into which *Hlj1^−/−^* BMDMs had been transplanted (Figure 6C). Similarly, macrophage-depleted *Hlj1^+/+^* mice receiving *Hlj1^−/−^* BMDM transplantation and LPS treatment showed lower serum levels of IL-12 and IFN-γ when compared to with those receiving *Hlj1^+/+^* BMDMs. Adoptive transfer of *Hlj1^+/+^* BMDMs into *Hlj1^−/−^* mice led to dramatically elevated mortality rates following an LPS challenge compared to *Hlj1^+/+^* mice transplanted with *Hlj1^+/+^* BMDMs (Figure 6D). In contrast, the survival of LPS-treated *Hlj1^+/^*^+^ mice was significantly improved when they were transplanted with *Hlj1^−/−^* BMDMs (Figure 6D). Combined, these results indicated that HLJ1 in macrophages is indispensable for maintaining IL-12–mediated IFN-γ production and contributing to septic death *in vivo*.

**Figure 6.**
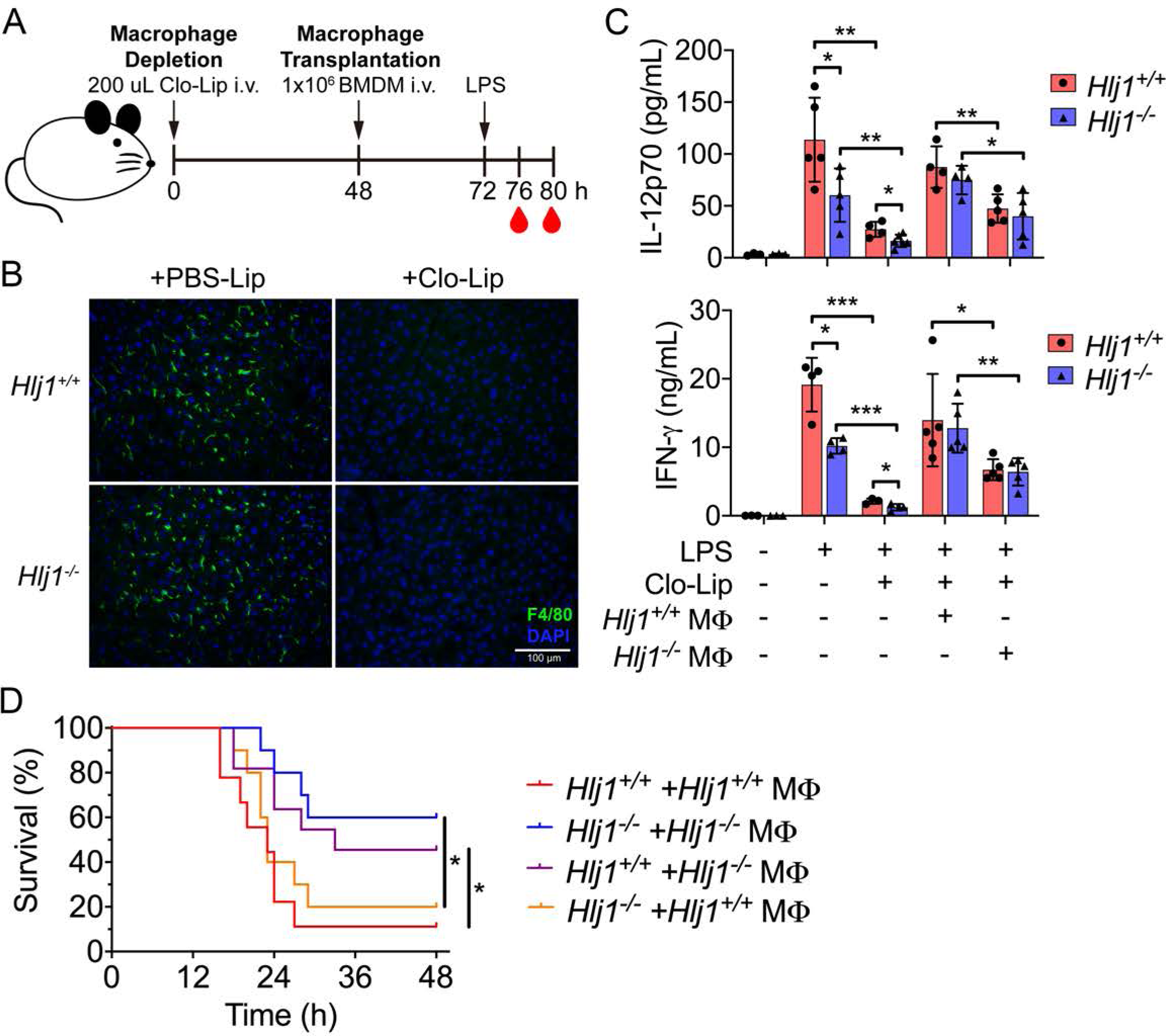
HLJ1 deletion in macrophages reduced serum levels of IL-12 and IFN-γ and mitigated septic death *in vivo*. (**A**) 200 μL clodronate liposomes (Clo-Lip) were administered intravenously to *Hlj1^+/+^* and *Hlj1^−/−^* mice to deplete their endogenous macrophages. After 48 h, the mice were intravenously injected with 1 × 10^6^ BMDMs isolated from *Hlj1^+/+^* or *Hlj1^−/−^* mice. After BMDM transplantation, *Hlj1^+/+^* and *Hlj1^−/−^* mice were administered with 20 mg/kg LPS and serum was collected at 4 h or 8 h for IL-12 or IFN-γ quantification, respectively. (**B**) Representative photographs of F4/80 immunofluorescence staining of liver sections from PBS liposome or clodronate liposome-injected *Hlj1^+/+^* and *Hlj1^−/−^* mice. The liver was fixed, dehydrated, embedded, cryosectioned into slices 8 μm thick, and incubated with anti-F4/80 antibodies to stain the mature macrophages (green). The scale bar represents 100 μm. (**C**) Mice transplanted with *Hlj1^+/+^* BMDMs (*Hlj1^+/+^* MΦ) or *Hlj1^−/−^* BMDMs (*Hlj1^−/−^* MΦ) were administered with LPS, and serum from n = 4–5 mice was analyzed for IL-12p70 and IFN-γ levels via ELISA. (**D**) Kaplan–Meier analysis of the overall survival of LPS-injected *Hlj1^+/+^* and *Hlj1^−/−^* mice transplanted with *Hlj1^+/+^* and *Hlj1^−/−^* BMDMs (n = 9–10 per group). For *Hlj1^+/+^* +*Hlj1^+/+^* MΦ vs *Hlj1^+/+^* +*Hlj1^−/−^* MΦ, *P* = 0.037. For *Hlj1^−/−^* +*Hlj1^−/−^* MΦ vs *Hlj1^−/−^* +*Hlj1^+/+^* MΦ, *P* = 0.036. Data presented are means ± SD. * *P* < 0.05, ** *P* < 0.01, *** *P* < 0.001. **Source data 1.** Data for graphs depicted in Figure 6C, D.

### HLJ1 helps IL-12p35 folding and heterodimeric IL-12p70 production

Macrophages are major innate immune cells responsible for IL-12 production in response to an endotoxin challenge, which leads to organ dysfunction and even septic shock. We isolated and differentiated BMDMs to investigate the underlying molecular mechanism of HLJ1-modulated IL-12 expression in macrophages. Up to 98% of BMDMs were obtained when differentiated under macrophage colony-stimulating factor (M-CSF) stimulation, demonstrating that the ability to differentiate did not differ between the genotypes (Figure 7– figure supplement 1A). When treated with LPS plus recombinant IFN-γ, *Hlj1^−/−^* BMDMs generated significantly less supernatant IL-12p70 in culture medium than *Hlj1^+/+^* BMDMs, but comparable quantities of IL-6 (Figure 7A, Figure 7– figure supplement 1B). Since intracellular IL-12 subunits were folded and assembled to allow secretion of biologically active heterodimeric IL-12, we assessed the ability to heterodimerization by using sandwich ELISA specifically detecting IL-12p70 heterodimer in BMDM cell lysate. Intracellular levels of heterodimeric IL-12p70 in *Hlj1^−/−^* BMDM were also lower than in *Hlj1^+/+^* BMDM (Figure 7B), suggesting that HLJ1 maintains the levels of IL-12p70 in macrophages.

**Figure 7.**
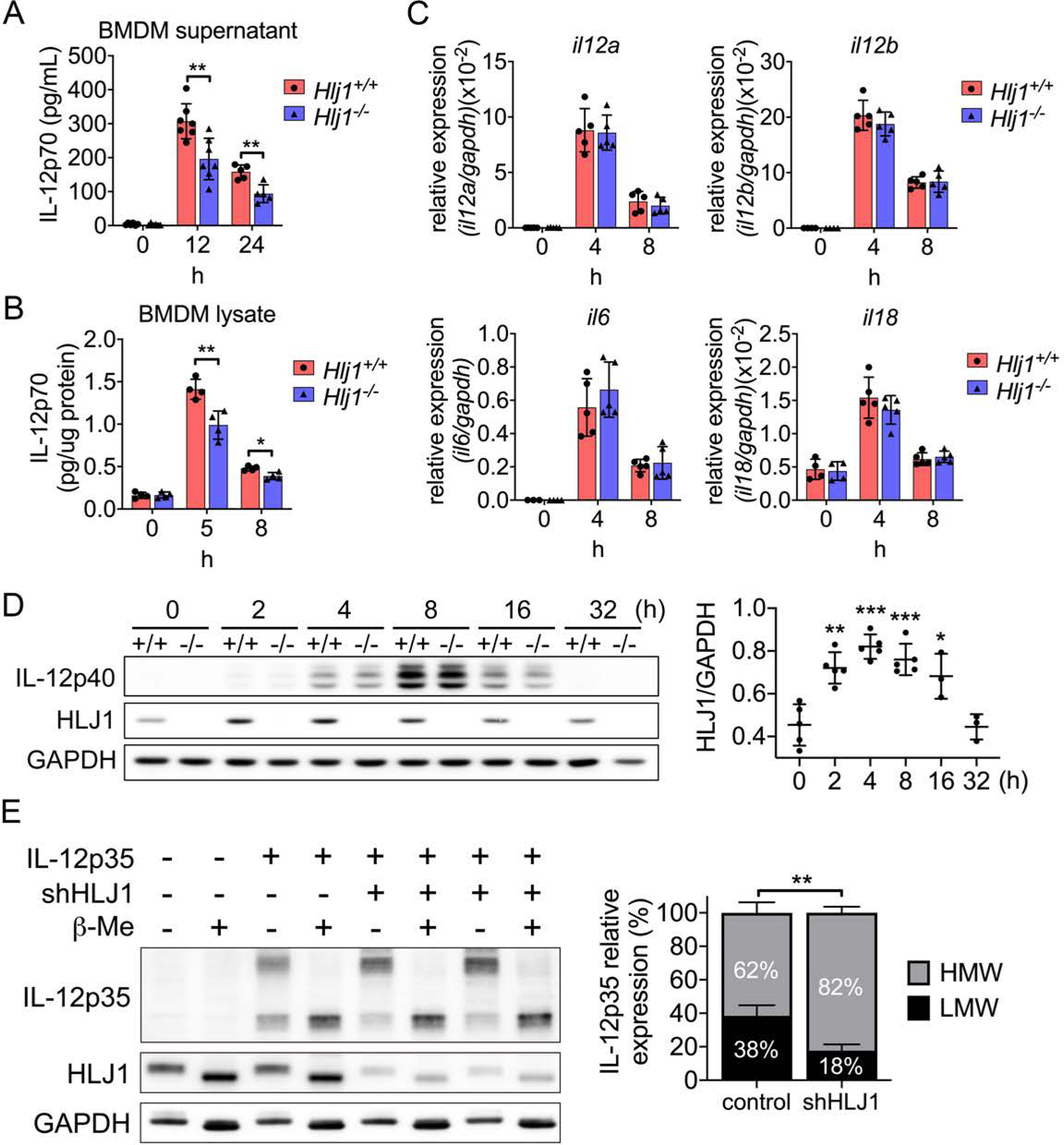
HLJ1 deletion leads to the accumulation of homodimeric IL-12p35 and reduced levels of heterodimeric IL-12p70. (**A**) BMDMs isolated from n = 6–7 *Hlj1^+/+^* and *Hlj1^−/−^* mice were treated with 10 ng/mL LPS and 20 ng/mL IFN-γ. Supernatant was collected at the indicated time points, and IL-12p70 was quantified via ELISA. 12 h, *P* = 0.032; 24 h, *P* = 0.025. (**B**) LPS/IFN-γ-treated BMDMs from n = 4–5 mice were lysed at the indicated time points and intracellular IL-12p70 was quantified via ELISA. 5 h, *P* = 0.006; 8 h, *P* = 0.012. (**C**) IL-12a, IL-12b, IL-6, and IL-18 expression was determined via qRT-PCR in LPS/IFN-γ–treated BMDMs isolated from n = 5 mice. (**D**) Intracellular IL-12p40 and HLJ1 expression levels were analyzed in LPS/IFN-γ-treated BMDMs isolated from *Hlj1^+/+^* (+/+) and *Hlj1^−/−^* (−/−) mice. Representative samples of n = 3–5 biological replicates are shown. GAPDH served as a loading control. In comparisons with the 0 h group (right panel): 2 h, *P* = 0.001; 4 h, *P* < 0.001; 8 h, *P* = < 0.001; 16 h, *P* = 0.02. (**E**) The influence of human HLJ1 knockdown on the redox state of human IL-12p35 was analyzed via nonreducing SDS-PAGE. 293T cells were (co-)transfected with the indicated IL-12p35 subunits and shRNA targeting HLJ1. The percentage of high molecular weight (HMW) and low molecular weight (LMW) IL-12p35 species in the presence or absence of shHLJ1 was quantified (right panel, n = 4 biological repeats for shHLJ1- and control-transfected cultures*; P* = 0.001). Where indicated, samples were treated with β-mercaptoethanol (β-Me) after cell lysis to provide a standard for completely reduced protein. GAPDH served as a loading control. Data presented are means ± SD. * *P* < 0.05, ** *P* < 0.01, *** *P* < 0.001. **Source data 1.** Data for graphs depicted in Figure 7A-E **Source data 2.** Original and labelled blots images of Figure 7D, E.

To understand how HLJ1 affects IL-12 expression, we performed qRT-PCR to evaluate the transcription levels of IL-12. Unexpectedly, we observed no difference in the mRNA levels of either IL-12p35 or IL-12p40 (subunits of IL-12p70) between *Hlj1^+/+^* and *Hlj1^−/−^* BMDMs upon treatment with either LPS/IFN-γ (Figure 7C) or LPS alone (Figure 7–figure supplement 1C), indicating that HLJ1 deletion has no effect on IL-12 transcriptional regulation. Transcriptional levels of the proinflammatory cytokines IL-6 and IL-18 also did not differ significantly between the two genotypes (Figure 7C and Figure 7– figure supplement 1C). We wondered if HLJ1 affects the quantity of IL-12 subunits, so we immunoblotted IL-12p40, a scaffold protein to maintain assembly-induced folding and secretion of IL-12, from LPS/IFN-γ-stimulated BMDM cell lysate, but we observed no significant difference between the genotypes (Figure 7D).

Because HSPs are proteins that can be induced upon cellular stress, we analyzed HLJ1 protein expression in response to LPS/IFN-γ cotreatment in BMDMs. HLJ1 protein levels increased in a time-dependent manner after LPS/IFN-γ stimulation in *Hlj1^+/+^* BMDMs, and decreased gradually from 8 h after the treatment (Figure 7D). However, hepatic HLJ1 expression remained unchanged in mouse liver, which is composed mainly of hepatocytes, after LPS injection (Figure 7 – figure supplement 2A and B), indicating that HLJ1 is induced in macrophages rather than in hepatocytes. Since serum levels of HSPs are reported to increase during sepsis, and analysis of survival outcomes of sepsis patients has revealed that increased mortality is associated with higher HSP serum levels, we postulated that HLJ1 may be secreted into the blood, and therefore quantified HLJ1 protein in serum from LPS-challenged mice. Indeed, we detected abundant HLJ1 in the serum of *Hlj1^+/+^* but not *Hlj1^−/−^* mice (Figure 7– figure supplement 2C). After the mice had received the LPS challenge, their serum levels of IL-12p70 were positively correlated to the amount of HLJ1 in their serum (Figure 7– figure supplement 2C).

Since chaperones have recently emerged as mediators of IL-12 family protein folding, assembly, and degradation, we performed simultaneous human IL-12p35 overexpression and human HLJ1 knockdown in 293T cells (an established model cell line for studying IL-12 family assembly) to assess the role of HLJ1 in regulating IL-12 biosynthesis. β-mercaptoethanol treatment breaks disulfide bonds and thus high-molecular-weight (HMW) IL-12p35 became completely reduced LMW IL-12p35 monomers, suggesting the HMW IL-12p35 was homodimer with intermolecular disulfide bridges between monomers (Figure 7E). Interestingly, HLJ1 knockdown shifted the HMW to LMW ratio of IL-12p35 to the dimeric species, suggesting that HLJ1 is able to prevent non-native homodimeric IL-12p35 accumulation and helps to reduce the disulfide bonds of homodimeric IL-12p35 to form LMW monomers (Figure 7E). These results indicated that LPS-inducible HLJ1 plays an important role in the regulation of IL-12p35 folding and the maintenance of IL-12p70 heterodimerization in endotoxin-stimulated primary macrophages. HLJ1-mediated production of IL-12p70 in macrophages has a substantial impact on the production of IFN-γ from NK cells and causes a persistent cytokine storm, contributing to endotoxin-induced lethality (Figure 8).

**Figure 8.**
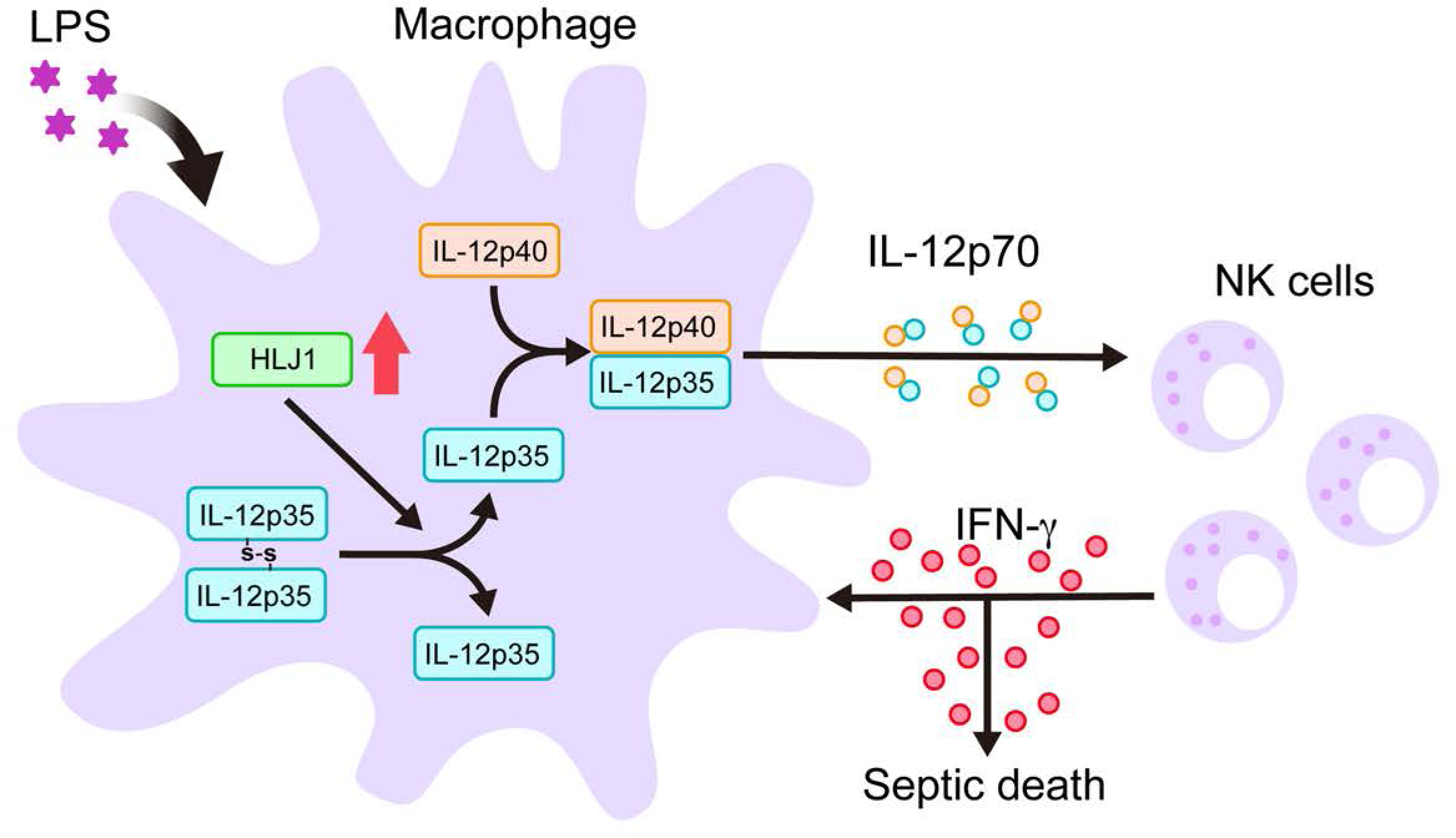
Schematic diagram depicting how HLJ1 controls IL-12 biosynthesis and the IFN-γ cytokine storm. The quantity of HLJ1 protein increases when macrophages are treated with LPS. The HLJ1 induced by LPS helps to reduce the quantity of HMW IL-12p35 homodimers, which becomes IL-12p35 monomers. Properly folded and assembled IL-12p70, which is composed of IL-12p35 and IL-12p40, is released by macrophages into the circulation and thereby stimulates NK cells. Eventually, activated NK cells release IFN-γ in sufficient quantities to lead to death via sepsis

## Discussion

Sepsis is a growing health problem, with multiple organ dysfunction syndrome developing in 30% of patients with sepsis. Liver dysfunction and failure, which are particularly serious complications of sepsis, contribute directly to disease progression and death (Canabal & Kramer, 2008). Although it is well known that the liver is one of the major organs that manages endotoxemia and is responsible for systemic immune modulation (Dhainaut et al., 2001; Siore et al., 2005), our knowledge about the transcriptional profiles of individual hepatic immune cells under the systemic inflammatory response of sepsis is limited. We used scRNA-seq to characterize the cellular landscapes of liver nonparenchymal cells in a mouse model of sterile sepsis. Enrichment analysis of differentially expressed genes demonstrated that HLJ1 deletion led to altered gene signatures in the IFN-γ–activating pathway of hepatic macrophages and dendritic cells in LPS-treated mice (Figure 1F and G). Consistent with previous studies, our data show that hepatic NK cells produce substantial quantities of IFN-γ in response to LPS stimulation, which has been reported to contribute to cytotoxicity against microbe-infected cells (Ami et al., 2002; Seki et al., 2000). In addition, hepatic T cells produce significantly greater quantities of IFN-γ, while other nonparenchymal cells generate trace amounts. B cells also produce IFN-γ, but to a lesser extent (Figure 2B and C). We subsequently found that HLJ1 deletion led to decreased IFN-γ signaling in NK cells but not in T and B cells, which also released IFN-γ in response to LPS challenge. These phenomena were not restricted to the liver: we found that splenic NK cells also produced less IFN-γ in *Hlj1^−/−^* mice (Figure 4A). In line with a recent scRNA-seq study showing that MHC genes are downregulated in monocytes in PBMC samples from patients with sepsis relative to those from control patients (Wen et al., 2020), our enrichment analysis showed that signaling pathway of MHC-I–dependent antigen presentation was altered and HLJ1 deletion had a substantial impact on it (Figure 1F).

Given that MHC is regarded as a potential biomarker and therapeutic target for sepsis (Venet et al., 2013), the role of HLJ1 in MHC-dependent antigen presentation warrants further investigation. Our results provide insights into intrahepatic transcriptional profiles and cellular landscapes under endotoxin stresses and underscore a novel role for HLJ1 in the immune modulation during endotoxic sepsis.

In the LPS mouse model, we demonstrated that endotoxin-induced death was attributable mainly to HLJ1-mediated production of IL-12 and IFN-γ–related cytokine storm *in vivo*. Here, mice received an intraperitoneal injection of 20 mg/kg LPS to induce endotoxic shock and systemic inflammation, which resembles sepsis-induced SIRS and severe endotoxemia (Silva et al., 2019). We demonstrated the effect of HLJ1 deletion on IFN-γ production by using high dose of LPS because mice are rather insensitive to LPS (Cauwels et al., 2013). Nonetheless, non-lethal low dose of LPS is sufficient to cause systemic inflammation as serum inflammatory cytokines such as TNF, IL-6, IFN-γ and IL-10 rapidly increase (Seemann et al., 2017). In order to understand the effect of non-lethal lower dose of LPS, we also treated both mice with 4 mg/kg of LPS which induced moderate endotoxemia (Kunze et al., 2019; Malgorzata-Miller et al., 2016). As it turned out, 4 mg/kg LPS injection led to lower serum levels of IFN-γ in *Hlj1^-/-^* mice comparing to wild-type mice (Figure 3D), suggesting the effect of HLJ1 on augmenting IFN-γ secretion can be also found during moderate endotoxemia.

In agreement with previous studies showing that IFN-γ acts on APCs to augment IL-12 transcription in a positive feedback loop (Grohmann et al., 2001; Ma et al., 1996), HLJ1 deficiency led to reduced IFN-γ levels and therefore reduced transcription of IL-12 in mice injected with LPS. Similarly, IFN-γ–stimulating pathways were altered in HLJ1-deleted macrophages and dendritic cells relative to wild-type cells in our scRNA-seq data. Bioactive IL-12 is a disulfide-bridged heterodimeric glycoprotein that consists of an α subunit (IL-12p35) and a β subunit (IL-12p40) (Yoon et al., 2000). After HLJ1 deletion, LPS-treated BMDMs exhibited no alteration in their mRNA levels of either IL-12p35 or IL-12p40, indicating that HLJ1 does not regulate IL-12 transcription directly. Nonetheless, we concluded that HLJ1 controlled the heterodimerization and maintained levels of the biologically active IL-12p70 protein, since HLJ1 deletion resulted in reduced intracellular and supernatant levels of heterodimeric IL-12p70 in LPS/IFN-γ–stimulated BMDMs. Recent studies have underscored the important role of chaperones in the maintenance of the assembly and folding of proteins in the IL-12 family. Meier et al. showed how chaperones regulate and control the assembly of heterodimeric IL-23, a member of the IL-12 family, through sequential checkpoints (Meier et al., 2019). Assembly-induced folding is thought to be a general mechanism in the biosynthesis of proteins in the IL-12 family. An IL-23 α-subunit is incompletely folded until a corresponding β-subunit bonds with it. Similarly, IL-12p35 folds inadequately and forms non-native disulfide bonds in the absence of IL-12p40. The misfolding of IL-12p35 is inhibited by IL-12p40, which gives rise to the native and bioactive IL-12 heterodimer (Reitberger et al., 2017). Chaperones also modulate quality-control checkpoints by recognizing incomplete folding. ERdj5, another member of the DnaJ protein family, participates in the recognition and removal of non-native disulfides, as well as in the ERAD of misfolded proteins (Oka et al., 2013; Ushioda et al., 2008), and had recently been found to reduce the quantity of IL-12p35 with non-native disulfide bonds and may even decelerate IL-12p35 degradation (Reitberger et al., 2017). In this study, we demonstrated that HLJ1-mediated conversion of IL-12p35 homodimers to LMW monomers, which contributes directly to IL-12p70 heterodimerization, because the lack of HLJ1 reduces the quantity of intracellular heterodimeric IL-12p70 detected by sandwich ELISA without changing the levels of intracellular IL-12p40 which acts as a scaffold to maintain assembly-induced folding. Accordingly, more detailed *in vitro* studies are needed to address questions about whether HLJ1 acts as a reductase to reduce non-native disulfide bonds in IL-12p35 dimers and whether HLJ1-mediated monomer formation contributes to the ERAD of misfolded IL-12p35 proteins.

The functions of HSPs are not limited to the intracellular environment, since they can be released into the extracellular space and act as danger signals that activate innate immunity by binding to receptors on immune cells (Chen et al., 2007; Schmitt et al., 2007; Zininga et al., 2018). HSP60 circulating in the blood modulates the innate immune system via activation of TLR4 and TLR2, which serve as a signal for APCs such as macrophages and dendritic cells (Henderson & Pockley, 2010; Quintana & Cohen, 2011). Intriguingly, we found that macrophages produced HLJ1 in a time-dependent manner when stimulated with LPS/IFN-γ, and HLJ1 could be detected in the blood of LPS-treated wild-type mice. Notably, HSP40 confers virulence and colonization in murine sepsis models, and recombinant HSP40 induces proinflammatory cytokine production in macrophages via activation of the PI3K and JNK signaling pathways (Cui et al., 2017). In another study, where elevated levels of anti-HSP40 antibodies were found in the serum of patients with rheumatoid arthritis, it was shown that both *E. coli* HSP40 (DnaJ) and human HSP40 (Hdj1) stimulated the secretion of IL-10 and IL-6 by PBMCs. These cellular responses inhibited the division of CD4^+^ and CD8^+^ T cells in the patients (Tukaj et al., 2010). Consequently, further investigation of the virulent role of extracellular HLJ1 protein and its potential immunomodulatory effects on immune cells is warranted. On the other hand, HSP70 serum levels are elevated during sepsis, and analysis of survival outcome indicates that increased mortality in patients with sepsis is associated with higher HSP serum levels (Gelain et al., 2011). Given that we observed a positive correlation between the serum levels of HLJ1 and those of IL-12 in septic mice, the potential role of HLJ1 as a biomarker for the prediction of patient prognosis and survival is worthy of further investigation.

We found HLJ1 to be a modulator of IL-12 protein folding and biosynthesis. Without HLJ1, the HMW IL-12p35 dimer accumulated in macrophages and IL-12p70 production was disturbed, which led to a less severe IFN-γ cytokine storm. During the previous decade, therapeutic agents targeting the IL-12 family of proteins, especially IL-12 and IL-23, are tested preclinically or clinically for inflammatory diseases such as psoriasis, Crohn’s disease, rheumatoid arthritis, and multiple sclerosis (Teng et al., 2015). Ustekinumab, a therapeutic agent targeting IL-12p40, has been approved for treating psoriasis and psoriatic arthritis (Savage et al., 2015). In addition, cytokine storm originating from the IL-12/IFN-γ axis plays an important role in the modulation of immunity and the pathogenesis of COVID-19. Elevated serum IL-12 levels have been observed in patients with SARS-CoV-2 infection (C. Chen et al., 2020; N. Chen et al., 2020; Huang et al., 2020). SARS-CoV-2 infects the respiratory epithelial tissue and stimulates local innate immune cells to initiate cytokine storm via the generation of inflammatory cytokines such as IL-1, IL-6, IL-8, IL-12, TNF-α, and other chemokines, which then recruit further innate and adaptive immune cells, resulting in the sustained production of inflammatory cytokines like IL-2, TNF-α, and IFN-γ. Such an acute response can induce myelopoiesis and emergency granulopoiesis, which may further exacerbate lung and epithelial damage (Yang et al., 2021). In addition, the overproduction of systemic cytokines, particularly IL-2, IFN-γ, GM-CSF, and TNF-α, leads to anemia and can disturb coagulation and vascular hemostasis, leading to capillary leak syndrome, thrombosis, and disseminated intravascular coagulation (DIC) (Mangalmurti & Hunter, 2020). These events, triggered by severe infection, eventually result in acute respiratory syndrome, multi-organ failure, and death (N. Chen et al., 2020; Huang et al., 2020; Wang et al., 2020). Transplantation of angiotensin-converting enzyme 2–negative (ACE2^−^) mesenchymal stem cells has been proposed as an effective therapy against COVID-19, since it inhibits the secretion of IL-12, IFN-γ, and TNF-α (Chen et al., 2013; Leng et al., 2020). Here, we used the well-established technique of macrophage depletion and reconstitution to investigate the role of macrophages in various disease models, including LPS-induced endotoxemia in mice (Fu et al., 2020). Our results showed that the transplantation of HLJ1-deficient macrophages resulted in significantly reduced serum IL-12 and IFN-γ levels and even protected the mice from septic death, implying that novel HLJ1-targeting strategies could be considered as a potential therapy against a variety of inflammatory diseases associated with IL-12 overproduction and even SARS-CoV-2 infection. Whether HLJ1 can modulate COVID-19-mediated immunopathology and whether targeting HLJ1 would improve the outcomes of patients with COVID-19 is worthy of further investigation.

In conclusion, we have demonstrated the previously unknown role of HLJ1 as a regulator of IL-12 folding and biosynthesis in macrophages. Our data show unequivocally that upregulated HLJ1 induced by LPS enhances IL-12 heterodimerization and secretion in macrophages, contributing to endotoxin-induced mortality in mice via IFN-γ upregulation in NK cells. Furthermore, HLJ1 deletion in macrophages protected mice from the pathogenesis of IFN-γ cytokine storm. Given that HLJ1 plays an important role in mediating IL-12/IFN-γ axis−dependent sepsis severity, HLJ1 may serve as a molecular target for the development of novel antisepsis or immunomodulatory therapeutics.

## Materials and Methods

### Mice and animal experiments

The HLJ1 knockout (*Hlj1^−/−^*) mouse was generated by the gene targeting strategy to delete exon2 of *Hlj1* gene at the embryonic stage. The syngeneic genetic background of *Hlj1* was achieved by backcrossing to C57BL/6 mouse strain over ten generations. Mice deficient in HLJ1 exhibited elevated ER stress-mediated abnormal lipogenesis (paper submitted). All mice were hosted in a pathogen-free facility, maintained in filter-topped cages under standard 12 hr light-dark cycle, and fed standard rodent chow and water ad libitum. All experimental procedures performed were approved by the Institutional Animal Care and Use Committee (IACUC) with IACUC number 20120515 and 20201050 at National Taiwan University Medical College. Liver and spleen were excised into adequate size for immunoblotting and qRT-PCR analysis. At the indicated time points, blood was collected from submandibular and complete blood counts were analyzed with IDEXX ProCyte Dx™. Serum levels of HDL, LDL and ALT were analyzed with Cobas c111 (Roche).

### LPS administration and analysis

The intraperitoneal LPS injection strategy of endotoxemia mouse model was based on previous studies (Samie et al., 2018; Silva et al., 2020; Starr et al., 2010). In survival analysis, 6–8-week-old mice were intraperitoneally injected with low dose (10 mg/kg) or high dose (20 mg/kg) LPS (Sigma-Aldrich, L2630) from E. coli O111:B4. For organ pathology analysis, mice were injected with 20 mg/kg LPS and sacrificed at indicated time points. For cytokine neutralization, mice were i.p. injected with 100 μg anti-IL-12 (C17.8) or anti-IFN-γ (XMG1.2) 1hr before LPS administration.

### Single-cell RNA sequencing

*Hlj1^+/+^* and *Hlj1^−/−^* mice were intraperitoneally injected with 20 mg/kg LPS and sacrificed 8 hours post injection. Largest liver lobe of n=3 mice from same group were pooled together and grinded with gentleMACS™ Dissociators. Mouse hepatic parenchymal cells were digested and non-parenchymal cells were isolated by Mouse Liver Dissociation Kit (Miltenyi Biotec). Isolated cells passed through 40-um cell strainer were treated with Red Blood Cell Lysis Solution (Miltenyi Biotec) to lyse blood cells. To acquire cells with >90% viability, dead cells were removed with Dead Cell Removal Kit (Miltenyi Biotec). Briefly, cells were pelleted by centrifugation at 300g for 5 minutes and resuspended in buffer containing dead cell removal microbeads and incubated at room temperature for 15 minutes. Cell suspension was applied to the MS columns (MACS Cat# 130-042-201) and effluent containing live cells was collected. Live cells were then centrifuged at 300g for 5 minutes, resuspended in cold 0.04% BSA/PBS and counted with Bio-Rad’s TC20^TM^ automated cell counter. Live cells were prepared for scRNA-seq with the Chromium Single Cell 3’ Reagent Kits v3 (10X Genomics) according to the user guide. Briefly, ∼4800 cells were wrapped into each gel-beads in emulsion (GEMs, 10X Genomics) at a concentration of 500-1000 cells/μL on Single Cell 3’ Chips v3 (10x Genomics) by using 10X Chromium controller. For reverse transcription incubation, GEMs were transferred to Bio-Rad C1000 Touch™ Thermal Cycler, followed by post GEMs–RT Cleanup, cDNA Amplification according to the manufacturer’s instructions. Qubit™ dsDNA HS Assay Kit (Invitrogen) was used to quantified cDNA concentration and single cell transcriptome libraries were constructed using the 10x Chromium Single Cell 3’ Library (10x Genomics, v3 barcoding chemistry). Quality control was performed with Agilent Bioanalyzer High Sensitivity DNA kit (Agilent Technologies). Libraries were then purified, pooled and analyzed on Illumina NovaSeq 6000 S2 Sequencing System with 150-bp paired-end reads.

### scRNA-seq data analysis

More than two billion scRNA-seq reads were processed and analyzed with the default parameters of CellRanger single-cell software suite (v3.1.0). Base calling files generated by Illumina sequencer were demultiplexed according to the sample index. Sequences were then aligned to the mm10 reference for whole transcriptome analysis. Multiple samples were aggregated for the following analysis. Loupe browser and Seurat (v4.0.0) were used to perform visualization, quality control, normalization, scaling, PCA dimension reduction, clustering, and differential expression analysis(Stuart et al., 2019). Cells with UMI count of greater than 30,000, fewer than 500 or greater than 6,000 genes, and >10% of total expression from mitochondrial genes were excluded (Figure 1– figure supplement 2A). The remaining 11,651 cells were unsupervised clustered after aligning the top 12 dimensions and setting resolution to 0.5. The identity for each cluster was assigned according to marker genes for known non-parenchymal cell types in the mouse liver (Xiong et al., 2019; Zhao et al., 2019). Differentially expressed genes with absolute log fold change greater than 0.25 and P value less than 0.05 were used for pathway and network enrichment analysis on the Metacore website. The scRNAseq data was deposited to GEO as raw and processed files with accession number GSE182137.

### Quantitative real-time PCR

RNA was isolated with TRI reagent (Sigma) and reverse transcribed with High-Capacity cDNA Reverse Transcription Kits (Applied Biosystems) according to the manufacturer’s instructions. The cDNA was used for qRT-PCR analysis with Power SYBR Green PCR Master Mix (Applied Biosystems) performed on the ABI-7500 Fast Real Time PCR system. The mRNA expression level was normalized to the amount of GAPDH gene expression, and the values were calculated using the comparative threshold cycle method (2^−ΔCt^). Primer sequences were listed in table supplement 1.

### Multiplex bead array and ELISA

Serum sample were taken at 0, 4, 8, 18 after 20 mg/kg LPS injection. Cytokine array was performed with LEGENDplex Mouse inflammation Panel (Biolegend) according to manufacturer’s protocol. Data analysis was proceeded by using LEGENDplex^TM^ Data Analysis Software. Mouse serum levels of IFN-γ, IL-1α and IL-6 were quantified with Mouse IFN-γ ELISA MAX^TM^ Deluxe Set (Biolegend), Mouse IL-1α ELISA MAX^TM^ Deluxe Set (Biolegend) and Mouse IL-6 ELISA MAX^TM^ Deluxe Set (Biolegend) according to the manufacturer’s protocol. Mouse serum, BMDMs supernatant and intracellular IL-12p70 were quantified by using ELISA MAX^TM^ Deluxe Set Mouse IL-12 (p70) (Biolegend).

### Flow cytometry

Mouse spleens were grinded and connective tissues were removed to obtain splenocytes. Pelleted cells were resuspended by 2 mL ACK lysis buffer and incubated for 1 min to lyse red blood cells, followed by a wash with 10 mL FACS buffer (PBS containing 0.2% BSA). Pelleted splenocytes were resuspended in FACS containing anti-CD16/CD32 antibodies (Biolegend) to block the Fc receptors before staining. After incubation for 10 min at 4℃, 50 μL diluted antibodies including anti-CD19, CD3, CD4, CD8 and NK1.1 antibody (Biolegend) were added and incubated for 30 min in the dark. Stained cells were washed twice with FACS, resuspended in 300 μL FACS, and then analyzed by CytoFLEX flow cytometer (Beckman Coulter). For intracellular IFN-γ staining, mice were injected with 20 mg/kg LPS and after 2.5 hours splenocytes were isolated. Splenocytes were fixed and permeabilized with 100 μL BD cytofix/cytoperm (BD biosciences) for 20 min after surface staining. BD Perm/Wash™ Buffer (BD biosciences) was used to wash the cells. Pelleted cells were resuspended in BD Perm/Wash™ Buffer containing anti-IFN-γ antibodies (Biolegend) for 30 min and analyzed by CytoFLEX flow cytometer.

### Cell culture and transfections

293T cells were cultured in DMEM (Thermo Fisher Scientific) supplemented with 10% FBS (Merck Millipore), 100 units/mL penicillin, 100 µg/mL streptomycin, 0.25 µg/mL Amphotericin B (Thermo Fisher Scientific) and 2 mM L-glutamine (Thermo Fisher Scientific) at 37°C and 5% CO2. HLJ1-shRNA-containing vectors were obtained from the National RNAi Core Facility (Academia Sinica, Taiwan). Plasmid containing human IL-12p35 cDNAs were obtained from Origene. One day before transfection, 2×10^5^/well 293T cells were plated on 6-well dishes. Co-transfection was carried out in 6-well dishes using lipofectamine 2000 reagent (Thermo Fisher Scientific) by adding 1.2 μg of HLJ1-shRNA and 2 μg of IL-12p35 plasmid DNA according to the protocol of the manufacturer. For primary NK cell experiment, pelleted splenocytes isolated from mice spleen were resuspended in FACS for NK cells purification with Mouse NK Cell Purification Kit (Miltenyi Biotec) according to the. Purified NK cells were surface-stained with anti-NK1.1 antibodies (Biolegend) for flow cytometry analysis, or were cultured in RPMI 1640 medium (Thermo Fisher Scientific) containing 10% FBS, 100 units/mL penicillin, 100 µg/mL streptomycin, 0.25 µg/mL Amphotericin B and 2 mM L-glutamine and treated with 10 ng/mL recombinant IL-12p70 (Peprotech) for 24 hours for supernatant IFN-γ analysis.

### BMDM isolation and activation

Bone marrow was flushed from murine femur and tibia and isolated cells were cultured in complete DMEM (Thermo Fisher Scientific) supplemented with 10 ng/mL M-CSF (Peprotech) (Trouplin et al., 2013). Three days after treatment, culture medium was replaced with fresh complete DMEM with 10 ng/mL M-CSF. At day 7, 10^6^/well differentiated macrophages were seeded into 6-well dishes. The next day BMDMs were classically activated with LPS (10 ng/mL) plus recombinant IFN-γ (Peprotech) (20 ng/mL), or stimulated with LPS alone (100 ng/mL).

### Macrophage depletion and reconstitution

Liposome-based macrophage depletion followed by BMDM reconstitution was described previously (Au -Weisser et al., 2012). Liposome-encapsulated clodronate (Liposoma) (100 μL per 10 g body weight) was i.v. administrated to deplete macrophages 3 days before LPS challenge. Adoptive transfer of macrophages was performed by i.v. injecting 1×10^6^ BMDMs 2 days after macrophage depletion.

### Immunoblotting experiments

Cells and mouse tissues were lysed in M-PER or T-PER Tissue Protein Extraction Reagent (Thermo Fisher Scientific) containing additional 1X PhosStop (Sigma) phosphatase inhibitor and 1X protease inhibitor cocktail (Sigma) and protein was extracted according to manufacturer’s protocol. For non-reducing SDS-PAGE gels, 20 mM N-Ethylmaleimide (NEM) (Sigma) was added to the lysis buffer. Samples were supplemented with 0.25 volumes of 4X sample buffer containing either 2-Me for reducing SDS-PAGE or 80 mM NEM for non-reducing SDS-PAGE. Samples were run on 10% SDS-PAGE gels, transferred to PVDF membranes, and blotted with anti-IL-12p35 (Abcam, Cat. ab133751), anti-IL-12p40 (Biolegend, Cat. 505310), anti-HLJ1 (Proteintech, Cat. 13064-1-AP) or anti-GAPDH (Proteintech, Cat. 60004-1-Ig) antibodies. Image J Software was used to semi-quantify the Western blotting results.

### Statistical analysis

Statistical analysis was performed by using the two-tailed, unpaired Student’s t-test with equal variance assumed. Log-rank Mantel-Cox test was used to compare survival curve. Correlation between serum HLJ1 and IL-12 was analyzed by using nonparametric Spearman correlation test. In scRNA-seq, differentially expressed genes of specific cell types were identified by using a Wilcoxon Rank Sum test. Differences were considered statistically significant when p<0.05.

## Data Availability

The raw and processed 10x single-cell sequencing data generated in this study have been deposited in the NCBI Gene Expression Omnibus (GEO) database under accession code GSE182137.

## Acknowledgements

We thank Pharmacogenomics Lab (TR6) and Center for Genomic and Precision Medicine, National Taiwan University for scRNA-seq technical support, and Hung-Wen Chen and Chia-I Lin (National Taiwan University) for technical support. This work was supported by grants MOST110-2314-B-002-269 (KYS) and MOST105-2628-B-002-051-MY3 (KYS) from Ministry of Science and Technology, Taiwan.

## Competing Interest Statement

All the authors declare that they have no competing interest.

---

**Figure supplement 1.**
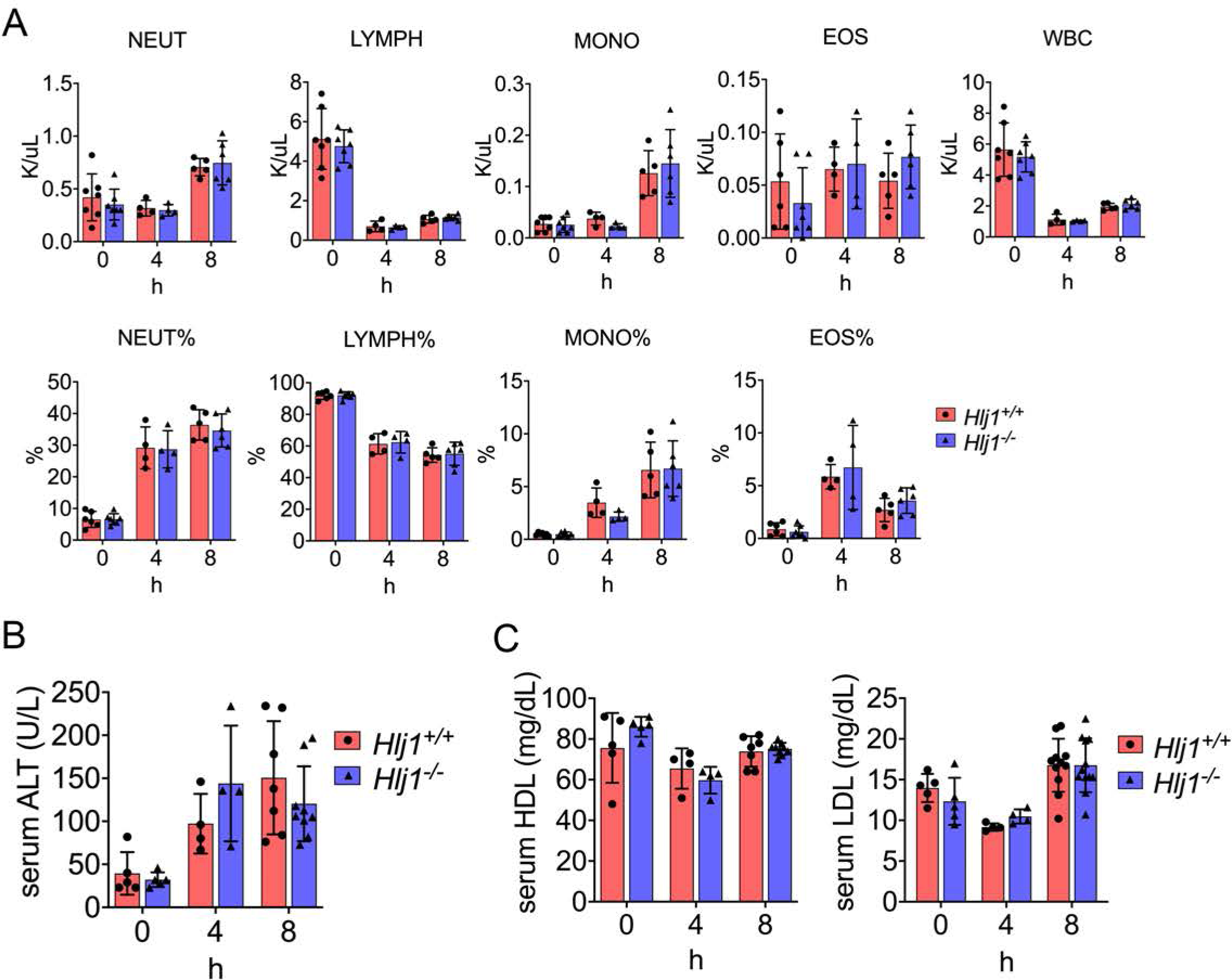
CBC counts, serum ALT, HDL and LDL levels of *Hlj1^+/+^* and *Hlj1^−/−^* mice. **(A)** 6–8 weeks *Hlj1^+/+^* and *Hlj1^−/−^* mice were injected with LPS of high dose (20 mg/kg). After 4 and 8 h, blood was collected for analyzing percentage and counts of neutrophils (NEUT), lymphocytes (LYMPH), monocytes (MONO), eosinophils (EOS) and white blood cells (WBC) (n = 5–7 mice). **(B)** Serum from n = 5-9 LPS–injected mice was analyzed for ALT levels. **(C)** Serum from n = 5-10 LPS– injected mice was analyzed for HDL and LDL levels. Data are mean ± SD. **Figure supplement 1-Source data 1.** Data for graphs depicted in Figure 1–figure supplement 1A-C.

**Figure supplement 2.**
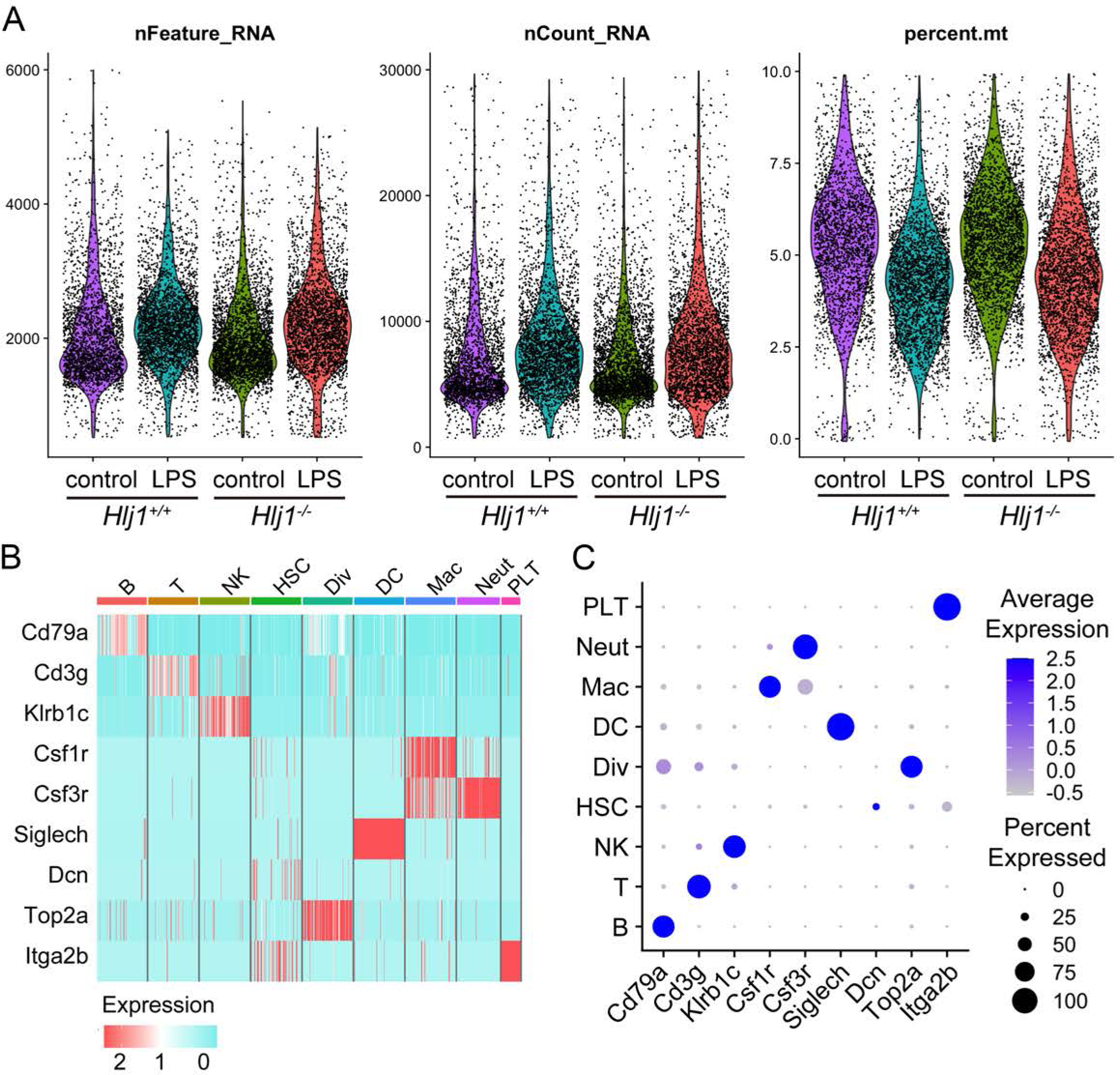
Quality control of scRNA–seq data. **(A)** Cells with unique molecular identifier (UMI) count of greater than 30,000, fewer than 500 or greater than 6,000 genes, and >10% of total expression from mitochondrial genes were excluded. **(B)** Dot plot and **(C)** heat map of known marker genes expression in each cell clusters.

**Figure supplement 1.**
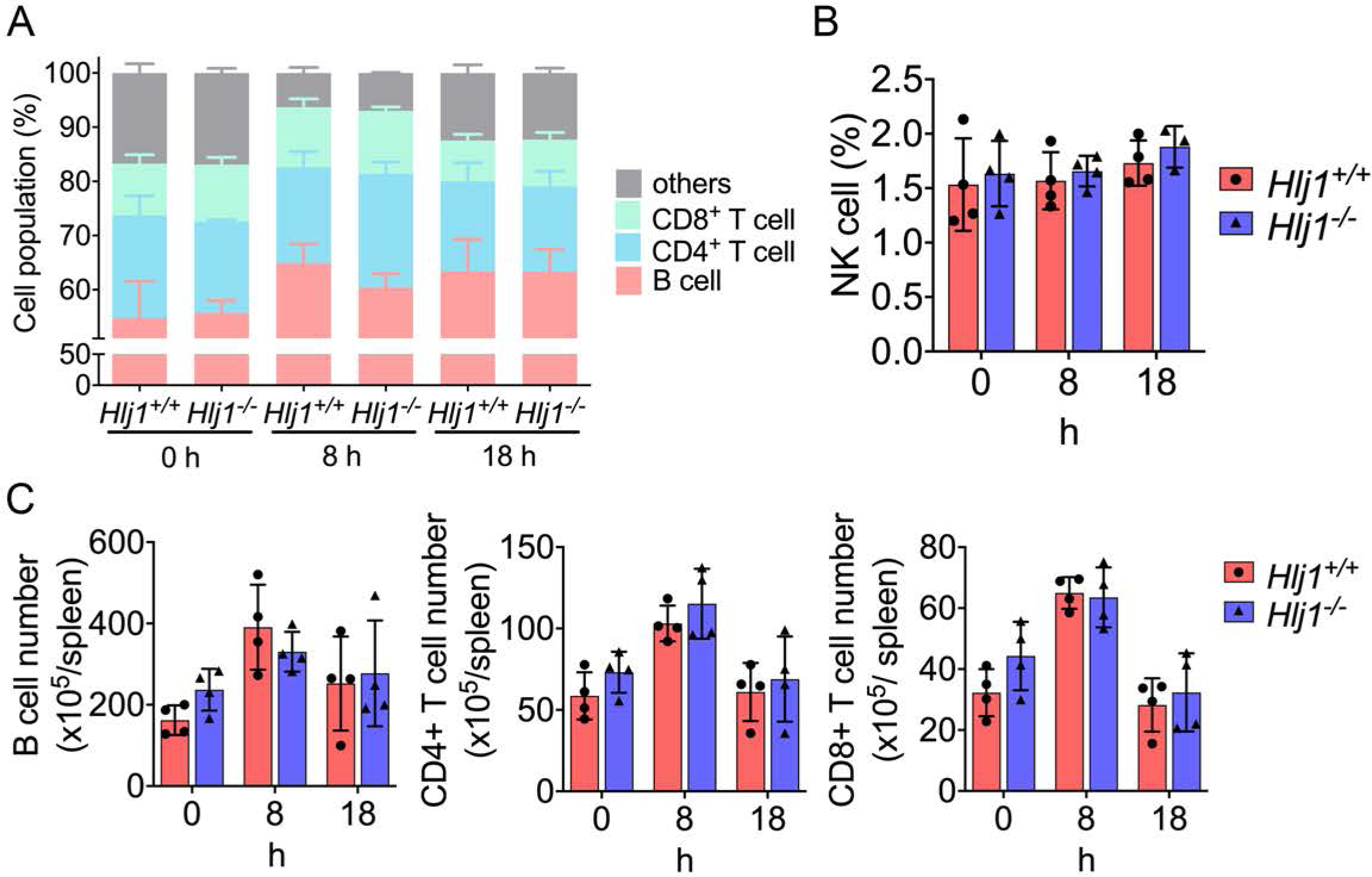
Splenic immune cell population identification of *Hlj1^+/+^* and *Hlj1^−/−^* mice. **(A)** CD4^+^, CD8^+^ T cell and B cell population and **(B)** NK cell population are presented as percentage. Splenic immune cells were isolated from LPS-treated *Hlj1^+/+^* and *Hlj1^−/−^* mice and were identified with surface markers of B, CD4^+^ T and CD8^+^ T cells by flow cytometry. **(C)** Immune cells population are presented as total number. **Figure supplement 1– Source data 1.** Data for graphs depicted in Figure 4– figure supplement 1A-C.

**Figure supplement 1.**
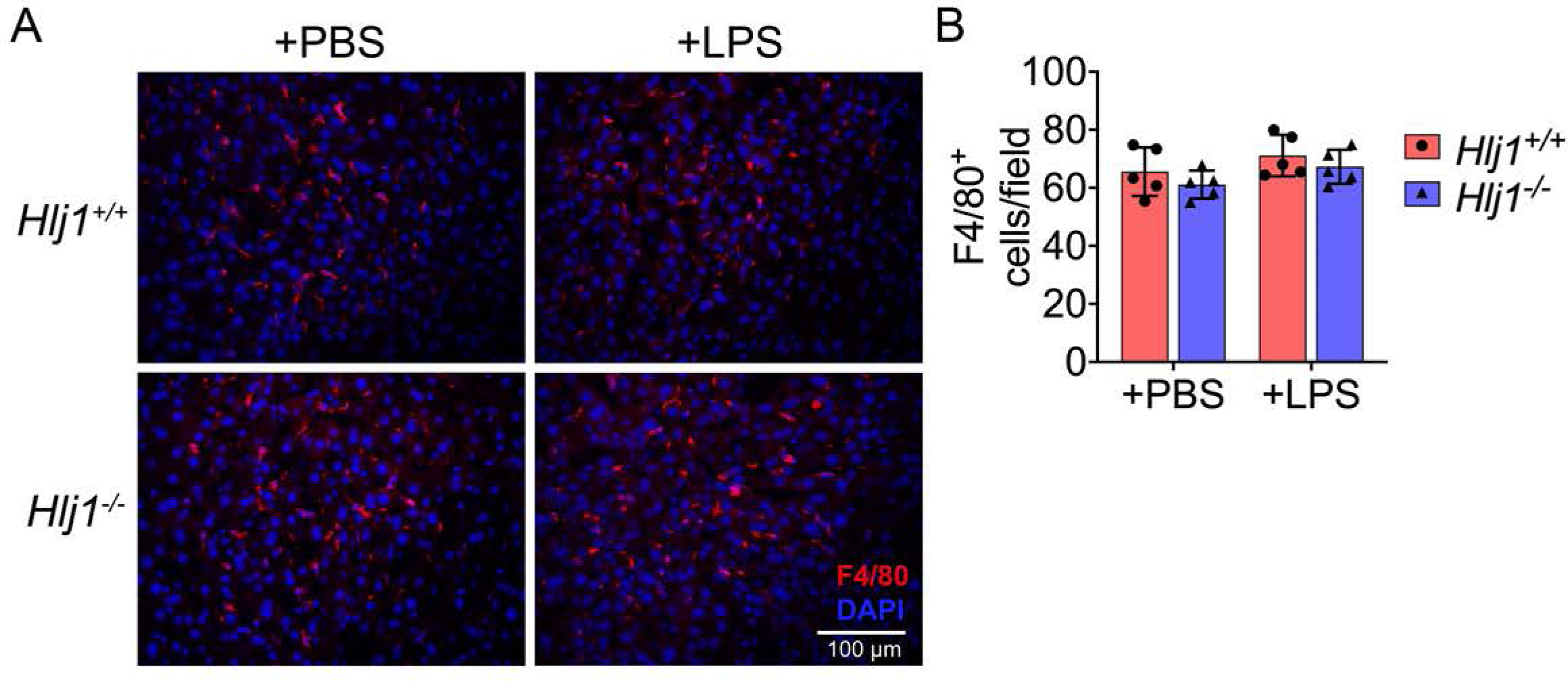
Quantification of liver-resident macrophages in LPS-challenged mice. **(A)** Representative photographs of F4/80 immunofluorescence staining from liver sections of PBS or LPS–injected *Hlj1^+/+^* and *Hlj1^−/−^* mice. 8 h after LPS injection, mice were sacrificed and liver was fixed, dehydrated, embedded, cryosectioned into 8 μm thickness, and incubated with anti–F4/80 antibodies to stain mature macrophages (red). **(B)** Quantitation of F4/80^+^ macrophages. Positively stained cells were counted at 400× magnification in 6 fields from 3 sections/mouse and from 5 mice/group. Data are mean ± SD. **Figure supplement 1-Source data 1.** Data for graphs depicted in Figure 5–figure supplement 1B.

**Figure supplement 1.**
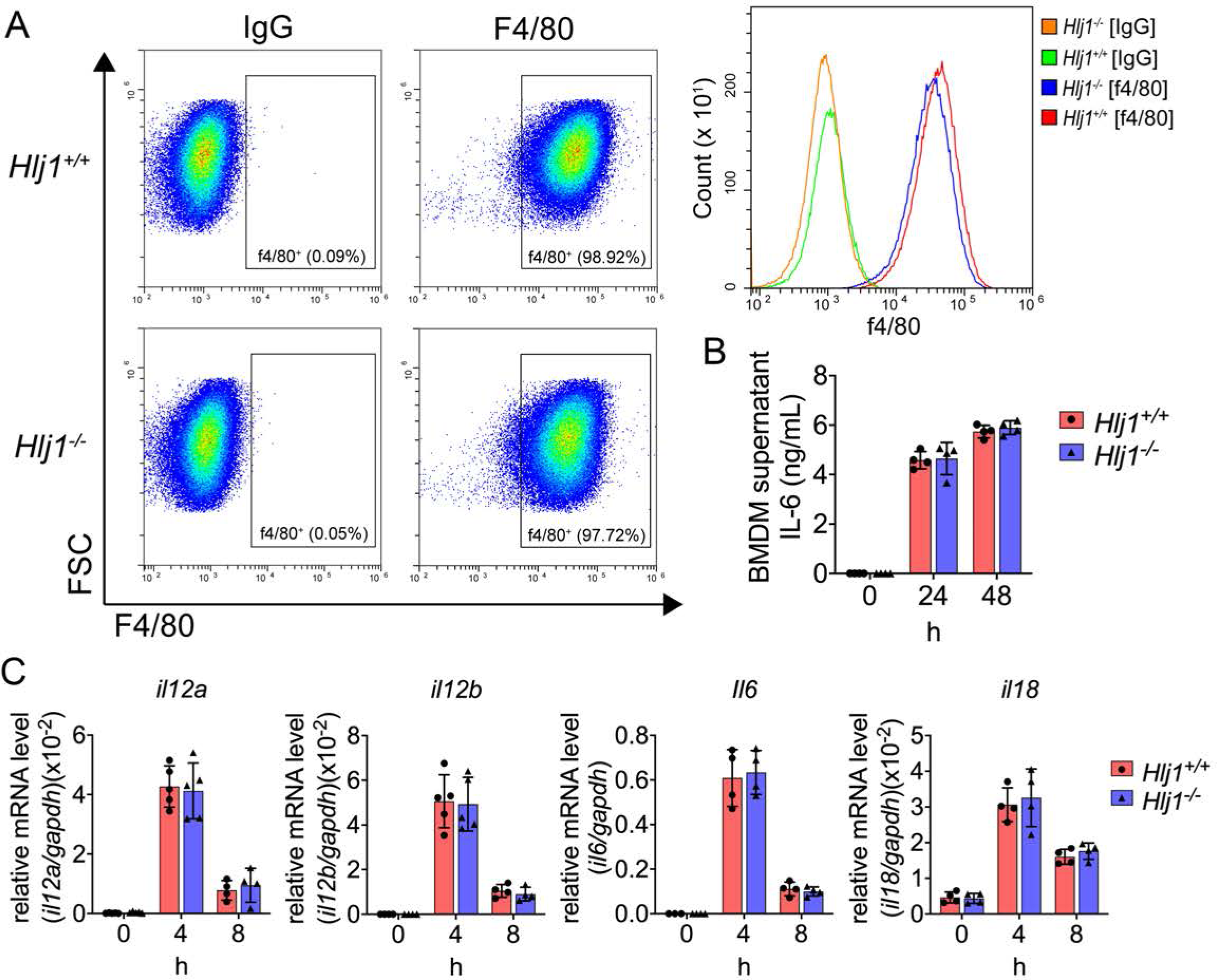
Transcriptional levels of proinflammatory cytokines in LPS– treated BMDMs. **(A)** F4/80^+^ BMDMs analyzed with flow cytometry. Bone marrow cells were isolated from *Hlj1^+/+^* and *Hlj1^−/−^* mouse and differentiated with M-CSF (10 ng/mL) for 7 days. **(B)** *Hlj1^+/+^* and *Hlj1^−/−^* BMDMs isolated from n = 4 mice were treated with 100 ng/mL LPS and supernatants were collected at the indicated time points and IL-6 was analyzed by ELISA. **(C)** Transcriptional levels of *il12a, il12b, il6 ad il18* in BMDMs isolated from n = 4–5 mice were quantified by qRT-PCR. Data are mean ± SD. **Figure supplement 1– Source data 1.** Data for graphs depicted in Figure 7–figure supplement 1B, C.

**Figure supplement 2.**
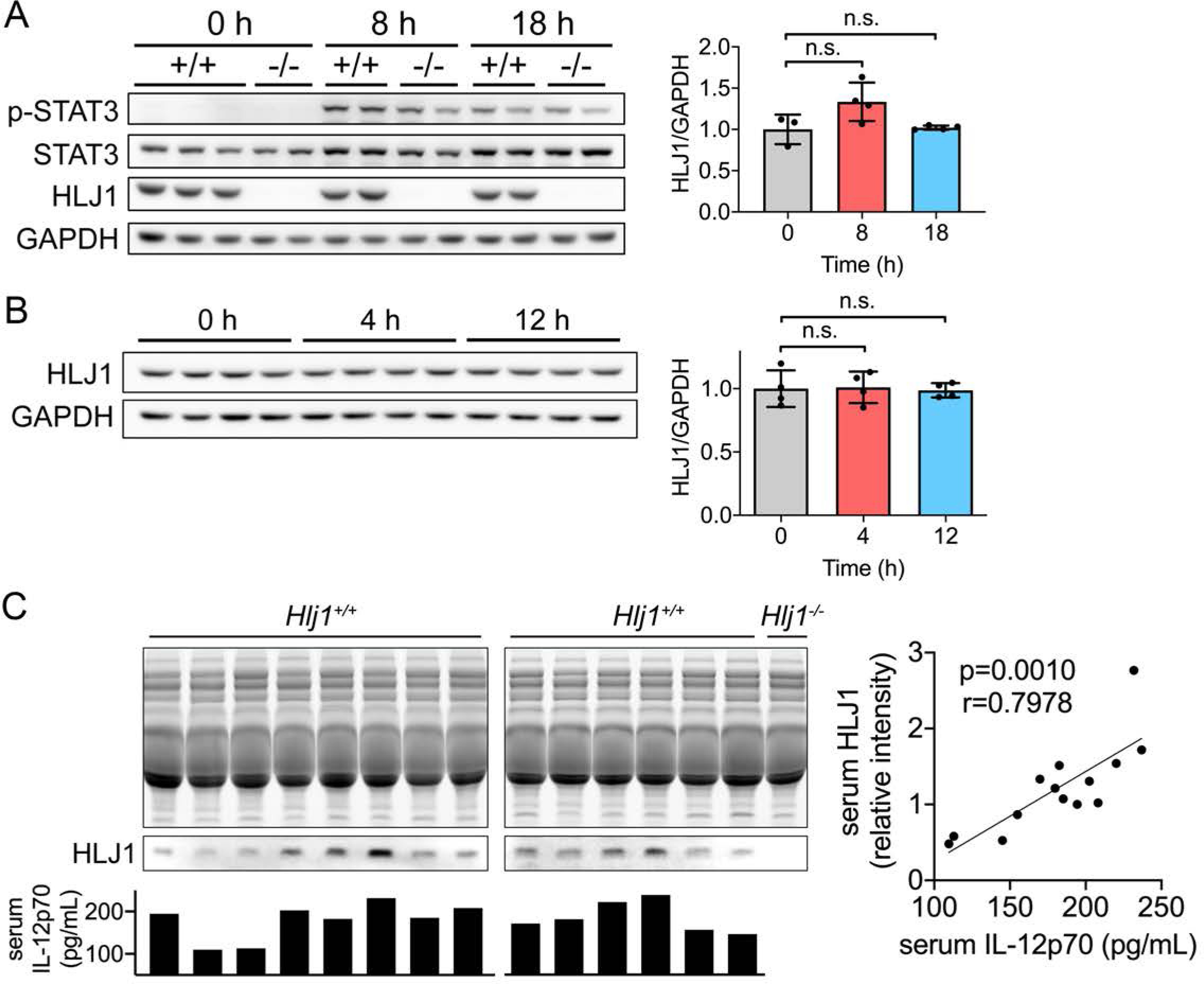
Expression levels of HLJ1 in the liver and serum from LPS– injected mice. **(A, B)** HLJ1 and STAT3 protein levels and phosphorylation levels in LPS-treated 6–8 weeks *Hlj1^+/+^* (+/+) and *Hlj1^−/−^* (*−/−*) mouse liver were determined. Representative samples are showed from n = 4 per group. Data are mean ± SD. **(C)** *Hlj1^+/+^* and *Hlj1^−/−^* mice were administrated with 20 mg/kg LPS and after 4 h serum was collected for analysis of IL-12p70 and HLJ1 expression by ELISA and Western blotting, respectively. For western blotting, 4 uL serum were used for each lane and total proteins in the acrylamide gel were stained as internal control. *P* value and r value were calculated from 14 samples by Spearman’s correlation coefficient testing. **Figure supplement 2– Source data 1.** Data for graphs depicted in Figure 7– figure supplement 2A-C. **Figure supplement 2– Source data 2.** Original and labelled blots images of Figure 7– figure supplement 2A-C.

## Notes

### Competing Interest Statement

The authors have declared no competing interest.

## References

1. Adhikari, N. K., Fowler, R. A., Bhagwanjee, S., & Rubenfeld, G. D. (2010). Critical care and the global burden of critical illness in adults. Lancet, 376(9749), 1339–1346. https://doi.org/10.1016/s0140-6736(10)60446-1

2. Ami, K., Kinoshita, M., Yamauchi, A., Nishikage, T., Habu, Y., Shinomiya, N., … Seki, S. (2002). IFN-γ Production from Liver Mononuclear Cells of Mice in Burn Injury As Well As in Postburn Bacterial Infection Models and the Therapeutic Effect of IL-18. The Journal of Immunology, 169(8), 4437. https://doi.org/10.4049/jimmunol.169.8.4437

3. Angus, D. C., & van der Poll, T. (2013). Severe Sepsis and Septic Shock. New England Journal of Medicine, 369(9), 840–851. https://doi.org/10.1056/NEJMra1208623

4. Au - Weisser, S. B., Au - van Rooijen, N., & Au - Sly, L. M. (2012). Depletion and Reconstitution of Macrophages in Mice. JoVE(66), e4105. https://doi.org/doi:10.3791/4105

5. Beutler, B., & Rietschel, E. T. (2003). Innate immune sensing and its roots: the story of endotoxin. Nature Reviews Immunology, 3(2), 169–176. https://doi.org/10.1038/nri1004

6. Canabal, J. M., & Kramer, D. J. (2008). Management of sepsis in patients with liver failure. Current Opinion in Critical Care, 14(2).

7. https://journals.lww.com/co-criticalcare/Fulltext/2008/04000/Management_of_sepsis_in_patients_with_liver.12.aspx

8. Cauwels, A., Vandendriessche, B., & Brouckaert, P. (2013). Of mice, men, and inflammation. Proceedings of the National Academy of Sciences, 110(34), E3150. https://doi.org/10.1073/pnas.1308333110

9. Chen, C., Zhang, X. R., Ju, Z. Y., & He, W. F. (2020). [Advances in the research of mechanism and related immunotherapy on the cytokine storm induced by coronavirus disease 2019]. Zhonghua Shao Shang Za Zhi, 36(6), 471–475. https://doi.org/10.3760/cma.j.cn501120-20200224-00088

10. Chen, H. W., Chen, H. Y., Wang, L. T., Wang, F. H., Fang, L. W., Lai, H. Y., … Hsu, S. C. (2013). Mesenchymal stem cells tune the development of monocyte-derived dendritic cells toward a myeloid-derived suppressive phenotype through growth-regulated oncogene chemokines. J Immunol, 190(10), 5065–5077. https://doi.org/10.4049/jimmunol.1202775

11. Chen, N., Zhou, M., Dong, X., Qu, J., Gong, F., Han, Y., … Zhang, L. (2020). Epidemiological and clinical characteristics of 99 cases of 2019 novel coronavirus pneumonia in Wuhan, China: a descriptive study. Lancet, 395(10223), 507-513. https://doi.org/10.1016/s0140-6736(20)30211-7

12. Chen, Y., Voegeli, T. S., Liu, P. P., Noble, E. G., & Currie, R. W. (2007). Heat shock paradox and a new role of heat shock proteins and their receptors as anti-inflammation targets. Inflamm Allergy Drug Targets, 6(2), 91–100. https://doi.org/10.2174/187152807780832274

13. Chousterman, B. G., Swirski, F. K., & Weber, G. F. (2017). Cytokine storm and sepsis disease pathogenesis. Seminars in Immunopathology, 39(5), 517–528. https://doi.org/10.1007/s00281-017-0639-8

14. Cui, J., Ma, C., Ye, G., Shi, Y., Xu, W., Zhong, L., … Wang, H. (2017). DnaJ (hsp40) of Streptococcus pneumoniae is involved in bacterial virulence and elicits a strong natural immune reaction via PI3K/JNK. Molecular Immunology, 83, 137–146. https://doi.org/10.1016/j.molimm.2017.01.021

15. Dhainaut, J. F., Marin, N., Mignon, A., & Vinsonneau, C. (2001). Hepatic response to sepsis: interaction between coagulation and inflammatory processes. Crit Care Med, 29(7 Suppl), S42–47. https://doi.org/10.1097/00003246-200107001-00016

16. Fajgenbaum, D. C., & June, C. H. (2020). Cytokine Storm. New England Journal of Medicine, 383(23), 2255–2273. https://doi.org/10.1056/NEJMra2026131

17. Fu, J., Zang, Y., Zhou, Y., Chen, C., Shao, S., Hu, M., … Zhang, T. (2020). A novel triptolide derivative ZT01 exerts anti-inflammatory effects by targeting TAK1 to prevent macrophage polarization into pro-inflammatory phenotype. Biomed Pharmacother, 126, 110084. https://doi.org/10.1016/j.biopha.2020.110084

19. Gelain, D. P., de Bittencourt Pasquali, M. A., C, M. C., Grunwald, M. S., Ritter, C., Tomasi, C. D., … Moreira, J. C. (2011). Serum heat shock protein 70 levels, oxidant status, and mortality in sepsis. Shock, 35(5), 466–470. https://doi.org/10.1097/SHK.0b013e31820fe704

20. Georgopoulos, C., & Welch, W. J. (1993). Role of the major heat shock proteins as molecular chaperones. Annu Rev Cell Biol, 9, 601–634. https://doi.org/10.1146/annurev.cb.09.110193.003125

21. Grohmann, U., Belladonna, M. L., Vacca, C., Bianchi, R., Fallarino, F., Orabona, C., … Puccetti, P. (2001). Positive Regulatory Role of IL-12 in Macrophages and Modulation by IFN-γ. The Journal of Immunology, 167(1), 221. https://doi.org/10.4049/jimmunol.167.1.221

22. Henderson, B., & Pockley, A. G. (2010). Molecular chaperones and protein-folding catalysts as intercellular signaling regulators in immunity and inflammation. J Leukoc Biol, 88(3), 445–462. https://doi.org/10.1189/jlb.1209779

23. Heymann, F., & Tacke, F. (2016). Immunology in the liver — from homeostasis to disease. Nature Reviews Gastroenterology & Hepatology, 13(2), 88–110. https://doi.org/10.1038/nrgastro.2015.200

24. Hotchkiss, R. S., Monneret, G., & Payen, D. (2013). Sepsis-induced immunosuppression: from cellular dysfunctions to immunotherapy. Nature Reviews Immunology, 13(12), 862–874. https://doi.org/10.1038/nri3552

25. Huang, C., Wang, Y., Li, X., Ren, L., Zhao, J., Hu, Y., … Cao, B. (2020). Clinical features of patients infected with 2019 novel coronavirus in Wuhan, China. Lancet, 395(10223), 497–506. https://doi.org/10.1016/s0140-6736(20)30183-5

26. Jirillo, E., Caccavo, D., Magrone, T., Piccigallo, E., Amati, L., Lembo, A., … Gumenscheimer, M. (2002). The role of the liver in the response to LPS: experimental and clinical findings. J Endotoxin Res, 8(5), 319–327. https://doi.org/10.1179/096805102125000641

27. Kumar, V. (2020). Toll-like receptors in sepsis-associated cytokine storm and their endogenous negative regulators as future immunomodulatory targets. International immunopharmacology, 89(Pt B), 107087–107087. https://doi.org/10.1016/j.intimp.2020.107087

28. Kunze, F. A., Bauer, M., Komuczki, J., Lanzinger, M., Gunasekera, K., Hopp, A. K., … Hottiger, M. O. (2019). ARTD1 in Myeloid Cells Controls the IL-12/18-IFN-gamma Axis in a Model of Sterile Sepsis, Chronic Bacterial Infection, and Cancer. J Immunol, 202(5), 1406–1416. https://doi.org/10.4049/jimmunol.1801107

29. Lechner, P., Buck, D., Sick, L., Hemmer, B., & Multhoff, G. (2018). Serum heat shock protein 70 levels as a biomarker for inflammatory processes in multiple sclerosis. Multiple sclerosis journal - experimental, translational and clinical, 4(2), 2055217318767192–2055217318767192. https://doi.org/10.1177/2055217318767192

30. Leng, Z., Zhu, R., Hou, W., Feng, Y., Yang, Y., Han, Q., … Zhao, R. C. (2020). Transplantation of ACE2(-) Mesenchymal Stem Cells Improves the Outcome of Patients with COVID-19 Pneumonia. Aging Dis, 11(2), 216–228. https://doi.org/10.14336/ad.2020.0228

31. Liu, Y., Zhou, J., Zhang, C., Fu, W., Xiao, X., Ruan, S., … Tang, M. (2014). HLJ1 is a novel biomarker for colorectal carcinoma progression and overall patient survival. Int J Clin Exp Pathol, 7(3), 969–977.

32. Ma, X., Chow, J. M., Gri, G., Carra, G., Gerosa, F., Wolf, S. F., … Trinchieri, G. (1996). The interleukin 12 p40 gene promoter is primed by interferon gamma in monocytic cells. J Exp Med, 183(1), 147–157. https://doi.org/10.1084/jem.183.1.147

33. Malgorzata-Miller, G., Heinbockel, L., Brandenburg, K., van der Meer, J. W., Netea, M. G., & Joosten, L. A. (2016). Bartonella quintana lipopolysaccharide (LPS): structure and characteristics of a potent TLR4 antagonist for in-vitro and in-vivo applications. Scientific reports, 6, 34221. https://doi.org/10.1038/srep34221

34. Mangalmurti, N., & Hunter, C. A. (2020). Cytokine Storms: Understanding COVID-19. Immunity, 53(1), 19–25. https://doi.org/10.1016/j.immuni.2020.06.017

35. Meier, S., Bohnacker, S., Klose, C. J., Lopez, A., Choe, C. A., Schmid, P. W. N., … Feige, M. J. (2019). The molecular basis of chaperone-mediated interleukin 23 assembly control. Nature Communications, 10(1), 4121. https://doi.org/10.1038/s41467-019-12006-x

36. Mencin, A., Kluwe, J., & Schwabe, R. F. (2009). Toll-like receptors as targets in chronic liver diseases. Gut, 58(5), 704. https://doi.org/10.1136/gut.2008.156307

37. Oka, O. B., Pringle, M. A., Schopp, I. M., Braakman, I., & Bulleid, N. J. (2013). ERdj5 is the ER reductase that catalyzes the removal of non-native disulfides and correct folding of the LDL receptor. Mol Cell, 50(6), 793–804. https://doi.org/10.1016/j.molcel.2013.05.014

38. Pierrakos, C., Velissaris, D., Bisdorff, M., Marshall, J. C., & Vincent, J. L. (2020). Biomarkers of sepsis: time for a reappraisal. *Critical care (London*, England), 24(1), 287. https://doi.org/10.1186/s13054-020-02993-5

39. Quintana, F. J., & Cohen, I. R. (2011). The HSP60 immune system network. Trends Immunol, 32(2), 89–95. https://doi.org/10.1016/j.it.2010.11.001

40. Reitberger, S., Haimerl, P., Aschenbrenner, I., Esser-von Bieren, J., & Feige, M. J. (2017). Assembly-induced folding regulates interleukin 12 biogenesis and secretion. The Journal of biological chemistry, 292(19), 8073–8081. https://doi.org/10.1074/jbc.M117.782284

41. Rivera, C. A., Adegboyega, P., van Rooijen, N., Tagalicud, A., Allman, M., & Wallace, M. (2007). Toll-like receptor-4 signaling and Kupffer cells play pivotal roles in the pathogenesis of non-alcoholic steatohepatitis. Journal of Hepatology, 47(4), 571–579. https://doi.org/10.1016/j.jhep.2007.04.019

42. Rosenzweig, R., Nillegoda, N. B., Mayer, M. P., & Bukau, B. (2019). The Hsp70 chaperone network. Nat Rev Mol Cell Biol, 20(11), 665–680. https://doi.org/10.1038/s41580-019-0133-3

43. Samie, M., Lim, J., Verschueren, E., Baughman, J. M., Peng, I., Wong, A., … Murthy, A. (2018). Selective autophagy of the adaptor TRIF regulates innate inflammatory signaling. Nat Immunol, 19(3), 246–254.https://doi.org/10.1038/s41590-017-0042-6

44. Savage, L. J., Wittmann, M., McGonagle, D., & Helliwell, P. S. (2015). Ustekinumab in the Treatment of Psoriasis and Psoriatic Arthritis. Rheumatology and therapy, 2(1), 1–16. https://doi.org/10.1007/s40744-015-0010-2

45. Schmitt, E., Gehrmann, M., Brunet, M., Multhoff, G., & Garrido, C. (2007). Intracellular and extracellular functions of heat shock proteins: repercussions in cancer therapy. J Leukoc Biol, 81(1), 15–27. https://doi.org/10.1189/jlb.0306167

46. Seemann, S., Zohles, F., & Lupp, A. (2017). Comprehensive comparison of three different animal models for systemic inflammation. Journal of Biomedical Science, 24(1), 60. https://doi.org/10.1186/s12929-017-0370-8

47. Seki, S., Habu, Y., Kawamura, T., Takeda, K., Dobashi, H., Ohkawa, T., & Hiraide, H. (2000). The liver as a crucial organ in the first line of host defense: the roles of Kupffer cells, natural killer (NK) cells and NK1.1 Ag+ T cells in T helper 1 immune responses. Immunol Rev, 174, 35–46. https://doi.org/10.1034/j.1600-0528.2002.017404.x

48. Silva, J. F., Olivon, V. C., Mestriner, F., Zanotto, C. Z., Ferreira, R. G., Ferreira, N. S., … Tostes, R. C. (2019). Acute Increase in O-GlcNAc Improves Survival in Mice With LPS-Induced Systemic Inflammatory Response Syndrome. Front Physiol, 10, 1614. https://doi.org/10.3389/fphys.2019.01614

49. Silva, J. F., Olivon, V. C., Mestriner, F. L. A. C., Zanotto, C. Z., Ferreira, R. G., Ferreira, N. S., … Tostes, R. C. (2020). Acute Increase in O-GlcNAc Improves Survival in Mice With LPS-Induced Systemic Inflammatory Response Syndrome [Original Research]. Frontiers in Physiology, 10(1614). https://doi.org/10.3389/fphys.2019.01614

50. Singer, M., Deutschman, C. S., Seymour, C. W., Shankar-Hari, M., Annane, D., Bauer, M., … Angus, D. C. (2016). The Third International Consensus Definitions for Sepsis and Septic Shock (Sepsis-3). JAMA, 315(8), 801–810. https://doi.org/10.1001/jama.2016.0287

51. Siore, A. M., Parker, R. E., Stecenko, A. A., Cuppels, C., McKean, M., Christman, B. W., … Brigham, K. L. (2005). Endotoxin-induced acute lung injury requires interaction with the liver. Am J Physiol Lung Cell Mol Physiol, 289(5), L769–776. https://doi.org/10.1152/ajplung.00137.2005

52. Starr, M. E., Ueda, J., Takahashi, H., Weiler, H., Esmon, C. T., Evers, B. M., & Saito, H. (2010). Age-dependent vulnerability to endotoxemia is associated with reduction of anticoagulant factors activated protein C and thrombomodulin. Blood, 115(23), 4886–4893. https://doi.org/10.1182/blood-2009-10-246678

53. Stuart, T., Butler, A., Hoffman, P., Hafemeister, C., Papalexi, E., Mauck, W. M., 3rd, … Satija, R. (2019). Comprehensive Integration of Single-Cell Data. Cell, 177(7), 1888-1902.e1821. https://doi.org/10.1016/j.cell.2019.05.031

54. Teng, M. W. L., Bowman, E. P., McElwee, J. J., Smyth, M. J., Casanova, J.-L., Cooper, A. M., & Cua, D. J. (2015). IL-12 and IL-23 cytokines: from discovery to targeted therapies for immune-mediated inflammatory diseases. Nature Medicine, 21(7), 719–729. https://doi.org/10.1038/nm.3895

55. Trouplin, V., Boucherit, N., Gorvel, L., Conti, F., Mottola, G., & Ghigo, E. (2013). Bone marrow-derived macrophage production. J Vis Exp(81), e50966. https://doi.org/10.3791/50966

56. Tsai, M. F., Wang, C. C., Chang, G. C., Chen, C. Y., Chen, H. Y., Cheng, C. L., … Yang, P. C. (2006). A new tumor suppressor DnaJ-like heat shock protein, HLJ1, and survival of patients with non-small-cell lung carcinoma. J Natl Cancer Inst, 98(12), 825-838. https://doi.org/10.1093/jnci/djj229

57. Tukaj, S., Kotlarz, A., Jozwik, A., Smolenska, Z., Bryl, E., Witkowski, J. M., & Lipinska, B. (2010). Hsp40 proteins modulate humoral and cellular immune response in rheumatoid arthritis patients. Cell stress & chaperones, 15(5), 555–566. https://doi.org/10.1007/s12192-010-0168-z

58. Ushioda, R., Hoseki, J., Araki, K., Jansen, G., Thomas, D. Y., & Nagata, K. (2008). ERdj5 is required as a disulfide reductase for degradation of misfolded proteins in the ER. Science, 321(5888), 569–572. https://doi.org/10.1126/science.1159293

59. Venet, F., Lukaszewicz, A.-C., Payen, D., Hotchkiss, R., & Monneret, G. (2013). Monitoring the immune response in sepsis: a rational approach to administration of immunoadjuvant therapies. Current opinion in immunology, 25(4), 477–483. https://doi.org/10.1016/j.coi.2013.05.006

60. Vincent, J. L., Marshall, J. C., Namendys-Silva, S. A., François, B., Martin-Loeches, I., Lipman, J., … Sakr, Y. (2014). Assessment of the worldwide burden of critical illness: the intensive care over nations (ICON) audit. Lancet Respir Med, 2(5), 380–386. https://doi.org/10.1016/s2213-2600(14)70061-x

61. Wang, C. C., Tsai, M. F., Hong, T. M., Chang, G. C., Chen, C. Y., Yang, W. M., … Yang, P. C. (2005). The transcriptional factor YY1 upregulates the novel invasion suppressor HLJ1 expression and inhibits cancer cell invasion. Oncogene, 24(25), 4081–4093. https://doi.org/10.1038/sj.onc.1208573

62. Wang, D., Hu, B., Hu, C., Zhu, F., Liu, X., Zhang, J., … Peng, Z. (2020). Clinical Characteristics of 138 Hospitalized Patients With 2019 Novel Coronavirus– Infected Pneumonia in Wuhan, China. JAMA, 323(11), 1061–1069. https://doi.org/10.1001/jama.2020.1585

63. Wen, M., Cai, G., Ye, J., Liu, X., Ding, H., & Zeng, H. (2020). Single-cell transcriptomics reveals the alteration of peripheral blood mononuclear cells driven by sepsis. Ann Transl Med, 8(4), 125. https://doi.org/10.21037/atm.2020.02.35

64. Wu, Y., Cui, J., Zhang, X., Gao, S., Ma, F., Yao, H., … Xu, W. (2017). Pneumococcal DnaJ modulates dendritic cell-mediated Th1 and Th17 immune responses through Toll-like receptor 4 signaling pathway. Immunobiology, 222(2), 384–393. https://doi.org/10.1016/j.imbio.2016.08.013

65. Xiong, X., Kuang, H., Ansari, S., Liu, T., Gong, J., Wang, S., … Lin, J.D. (2019). Landscape of Intercellular Crosstalk in Healthy and NASH Liver Revealed by Single-Cell Secretome Gene Analysis. Mol Cell, 75(3), 644–660.e645. https://doi.org/10.1016/j.molcel.2019.07.028

66. Yang, H., Young, D. W., Gusovsky, F., & Chow, J. C. (2000). Cellular events mediated by lipopolysaccharide-stimulated toll-like receptor 4. MD-2 is required for activation of mitogen-activated protein kinases and Elk-1. The Journal of biological chemistry, 275(27), 20861-20866. https://doi.org/10.1074/jbc.M002896200

67. Yang, L., Xie, X., Tu, Z., Fu, J., Xu, D., & Zhou, Y. (2021). The signal pathways and treatment of cytokine storm in COVID-19. Signal Transduction and Targeted Therapy, 6(1), 255. https://doi.org/10.1038/s41392-021-00679-0

68. Yoon, C., Johnston, S. C., Tang, J., Stahl, M., Tobin, J. F., & Somers, W. S. (2000). Charged residues dominate a unique interlocking topography in the heterodimeric cytokine interleukin-12 [https://doi.org/10.1093/emboj/19.14.3530]. The EMBO Journal, 19(14), 3530–3541. https://doi.org/10.1093/emboj/19.14.3530

69. Zhao, L., Jin, Y., Donahue, K., Tsui, M., Fish, M., Logan, C. Y., … Nusse, R. (2019). Tissue Repair in the Mouse Liver Following Acute Carbon Tetrachloride Depends on Injury-Induced Wnt/β-Catenin Signaling. Hepatology, 69(6), 2623–2635. https://doi.org/10.1002/hep.30563

70. Zhou, F., Yu, T., Du, R., Fan, G., Liu, Y., Liu, Z., … Cao, B. (2020). Clinical course and risk factors for mortality of adult inpatients with COVID-19 in Wuhan, China: a retrospective cohort study. Lancet, 395(10229), 1054–1062. https://doi.org/10.1016/s0140-6736(20)30566-3

71. Zininga, T., Ramatsui, L., & Shonhai, A. (2018). Heat Shock Proteins as Immunomodulants. Molecules, 23(11). https://doi.org/10.3390/molecules23112846

